# CryoVIA - An image analysis toolkit for the quantification of membrane structures from cryo-EM micrographs

**DOI:** 10.1101/2025.01.09.632183

**Authors:** Philipp Schönnenbeck, Benedikt Junglas, Carsten Sachse

**Affiliations:** Ernst-Ruska Centre for Microscopy and Spectroscopy with Electrons, ER-C-3/Structural Biology, Forschungszentrum Jülich, 52425 Jülich, Germany; Department of Biology, Heinrich Heine University, Universitätsstr. 1, 40225 Düsseldorf, Germany

**Keywords:** cryo-EM, image analysis, segmentation, membrane analysis, lipid vesicles, curvature, membrane thickness, vesicle shapes, membrane remodeling, ESCRT-III

## Abstract

Imaging of lipid structures and associated protein complexes using electron cryo-microscopy (cryo-EM) is a common visualization and structure determination technique. The quantitative analysis of the membrane structures, however, is not routine and time consuming in particular when large amounts of data are involved. Here, we introduce the automated image-processing software *cryo-vesicle image analyzer* (CryoVIA) that parametrizes lipid structures of large data sets from cryo-EM images. This toolkit combines segmentation, structure identification with methods to automatically perform a large-scale data analysis of local and global membrane properties such as bilayer thickness, size, and curvature including membrane shape classifications. We included analyses of exemplary data sets of different lipid compositions and protein-induced lipid changes through an ESCRT-III membrane remodeling protein. The toolkit opens new possibilities to systematically study structural properties of membrane structures and their modifications from cryo-EM images.

## Introduction

Cryo-EM is a powerful tool to study protein-membrane interactions as it provides high-resolution images of vitrified macromolecules under near-native conditions including biological membranes^1^. Isolated and soluble protein structures of biological macromolecules are commonly determined to high resolution enabling atomic model building^2^. A group of lipid-interacting proteins is capable of deforming lipid membranes and inducing shape changes on lipid membranes through their distinct binding mechanisms^3^. Archetypical examples are GTPase driven dynamin or bar-domain proteins that are involved in the cellular membrane-trafficking processes like endocytosis^4,5^. Another important group of membrane-remodeling proteins constitutes the family of endosomal sorting complexes required for transport III (ESCRT-III) that are topologically catalyzing budding reactions away from the cytosol. While structures of membrane-remodeling proteins can be determined in isolation, many of them require their biological substrate in order to polymerize^6,7^.

A series of membrane-remodeling protein structures have been determined in the presence of lipid membranes using cryo-EM^8–10^. For this purpose, Fourier-Bessel, single-particle or hybrid approaches of image processing approaches for helical reconstruction were employed to resolve the protein structures of the helical lattices formed on lipid membranes^11–13^. In the meantime, widely used single-particle image processing software suites have been adapted to work with helical protein assemblies^14,15^. These image processing suites primarily focus their refinement on protein structures. In the case when lipid membranes are tightly bound to the protein structures, lipid density can be observed alongside the protein density^16^. However, phospholipid bilayers alone also generate significant contrast suitable for detailed analysis of characteristic lipid features and membrane shapes^17^. Nevertheless, the localization of such membrane structures and the quantitative evaluation of those features is often performed interactively and remains a subjective as well as labor-intensive task. For the case of cellular electron tomograms, few automated membrane analysis tools have been put forward^18–20^ while they target to capture the larger scale cellular environment in three-dimensional image volumes.

Quantitative analysis of membrane structure from many micrographs is still a largely manual task as no suitable image processing tools are available. To this end, we developed a *cryo vesicle image analysis* (CryoVIA) tool to automate membrane structural analysis. The developed toolkit combines neural network-based image segmentation, structure identification with a series of analytical tools to estimate local and global structural properties including the classification of vesicular membrane shapes. We apply CryoVIA to different preparations of vesicles and lipid mixtures of exemplary data sets and reveal modifications to their structural properties.

## Results

### Principal workflow of CryoVIA

In order to generate a quantitative automated workflow suitable for analyzing biological membranes from cryo-micrographs, we created CryoVIA that requires electron micrographs as input and concludes with analytical data plots. The workflow is divided into three basic parts: neural-network based segmentation, membrane structure identification, and structure analysis (Fehler! Verweisquelle konnte nicht gefunden werden.). For the first part of the toolkit, the given micrographs are segmented to locate the membrane structures of interest that are subsequently used for further analysis. We applied segmentation using a pre-trained U-net with micrographs including user-provided membrane structures as input^21^ and found membranes across micrographs consistently labelled. By default, a pre-trained U-Net is used, while some use cases may require to re-train the U-Net on manually segmented micrographs.

### Identification of membrane features

**The second part of the toolkit refers to the membrane structure identification that will generate membrane entities based on the segmentation results from the first part. The results of the binary segmentation are used to identify independent instances of membrane structures within one micrograph by simply splitting the segmentation into independent connected components. This approach worked well when membrane structures are well separated in one micrograph, however, when the target structures are crowded and membranes overlap, they require additional separation routines. Overlapping membrane structures are located and re-connected by monitoring the continuity of local membrane structures (see** STAR methods for details). For presenting the principal functional features of CryoVIA, we, initially, use a test data set containing 356 cryo-micrographs of DOPC vesicles that were size-filtered by a 200 nm cutoff. As a result of the feature identification, the initially segmented skeletons are successfully separated by restraining subsequent angles into individual membrane structures (Fehler! Verweisquelle konnte nicht gefunden werden.**A**).

**For detailed and consistent analysis of the identified membrane structures, refinement of the initially detected pixel positions is essential to obtain accurate membrane contours located at the center of each membrane bilayer (Fehler! Verweisquelle konnte nicht gefunden werden.B). For this purpose, we convolve the image with a bilayer-like kernel and subsequently employ the Frangi filter**^22^ **(see** STAR methods for details). As a result of the membrane structure identification, the curated membrane contours are available for detailed analysis of the membrane structures. As an optional step, it is also possible to remove identified membranes outside of grid foil holes using a circle convolution (**Figure S1**). After identification of the foil hole edges, localized membrane structures outside of the holes can be effectively removed and the relevant membrane features are ready for subsequent in-depth data analysis. In summary, through a series of refinement and pruning steps, segmentation results are enhanced for the following analysis of structural features.

### Quantitative extraction of local and global membrane features

As the final part of the CryoVIA workflow, we perform structure analyses for each membrane contour using the information from the identified pixels. Measurements for each contoured pixel such as curvature, bilayer thickness, and distance to other membrane segments are obtained locally. Global statistics about the membrane structure such as length, area, diameter are directly derived from the membrane contours. Finally, shape classifications are employed for large-scale and intuitive comparisons of different experimental data sets.

In order to describe local shape changes of the membrane contours, we make use of the curvature descriptor. Curvature is defined as the reciprocal radius of a circle that best describes the local curve. The higher the curvature, the higher is the local shape changes of the membrane while a curvature of zero corresponds to a straight line. In addition, the direction of the curvature is calculated and represented by the sign of the curvature value. For closed membranes (e.g., vesicles), convex regions of the membrane have a positive curvature value while concave regions will assume a negative value. Based on a local measurement, it is not possible to distinguish between convex and concave curvature in membrane segments, as they require the topological information of the entire membrane entity. In the following analysis only completely closed membranes - vesicles - are considered and analyzed. The calculated curvature values are presented in two dimensions superimposed on the micrograph or as a function of the contour length (Fehler! Verweisquelle konnte nicht gefunden werden.**C/D**).

Another important local membrane parameter is the underlying bilayer thickness or leaflet separation reflecting the physical properties and chemical composition of the membrane^17^. For bilayer thickness estimation, the averaged local cross sections of the bilayer density are extracted and smoothened by a Gaussian filter until two distinct minima can be identified. The distance between these minima is estimated as the bilayer thickness (Fehler! Verweisquelle konnte nicht gefunden werden.**E/F**). Due to the inherent noise in cryo-EM images, the estimated thickness values along the contour of the vesicle are additionally smoothened by a Gaussian filter. The bilayer thickness is estimated for each contour pixel of the vesicle locally as well as averaged over one membrane structure, e.g., for one vesicle.

Global structural parameters such as diameter, circumference, and area of each identified membrane structure are directly extracted based on the membrane contours according to their geometric definitions (Fehler! Verweisquelle konnte nicht gefunden werden.**G**). Moreover, distances to other membrane structures are also determined, e.g., for cases when vesicles contain additional enclosed vesicles. Membranes and vesicles occur in a wide variety of shapes, therefore, classification or clustering into different classes is desirable for further analysis. As self-contained membrane structures can only assume a limited set of shapes^23^, they are classified by a small one-dimensional convolutional neural network using simulated reference shapes. Based on our most frequent shape observations, we designed the default classifier to distinguish seven shapes including sphere, prolate, tube, pear, stomatocyte, hourglass, and an elongated pear (Fehler! Verweisquelle konnte nicht gefunden werden.**H**). The shape classification makes use of the fact that the curvature along the contour of the vesicle has a distinct profile for each shape. To perform classification, the curvature contour of a given vesicle is length-normalized by resizing the curvature values to a predefined vesicle size, interpolating to a fixed number of data points and the values are shifted to start with the lowest value. The length-normalized values can then be used by the classifier. The default classifier identifies the test shapes with an accuracy of 98.6% after training on a set of 200 instances for each shape (Fehler! Verweisquelle konnte nicht gefunden werden.). In addition, it is possible to add custom shapes to classify specific vesicles after user definition using the provided GUI.

### Errors in curvature and bilayer thickness estimation

To estimate the precision of the local curvature estimation, we used another test data set of DOPC sonicated vesicles. We compared the local measurements with segmentation-based global parameters that are generally of higher reliability. For this purpose, we looked at 500 ideal vesicles of highest circularity and of diameters of >1000 Å. The circularity was estimated by global segmentation parameters 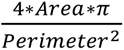 with 1 being a perfect circle and 0 being highly non-circular shapes, which we consider to be a reliable measurement. Next, we estimated the curvature locally and circle radii for every point of the vesicle using six different neighborhood sizes (50, 100, 200, 300, 400, and 500 Å, respectively) and subsequently calculated the difference between these locally estimated radii and the globally derived radii (Fehler! Verweisquelle konnte nicht gefunden werden.**A**). The errors obtained with large neighborhood sizes of 200, 300, 400, and 500 are small at 6.3, 3.4, 2.8, and 2.6 Å, respectively. When using small neighborhood sizes of 50 and 100 Å, the mean error in the curvature estimation was substantially higher at 44.8 Å and 18.1 Å, respectively, presumably due to the increase of noise over signal available for radii fitting. While small neighborhood sizes can detect radius changes more locally, they are more prone to overfitting in the presence of higher noise levels in the measurements. As a compromise between precision of the curvature estimation and locality of the detectable features, we chose a default neighborhood size of 200 Å.

To evaluate the error of the bilayer thickness estimation, we examined the forward difference of the estimated bilayer thickness values. For the same 500 most circular vesicles described above, the mean difference and standard deviation corresponded to 0.14 Å and 0.17 Å, respectively. Based on these considerations, bilayer thickness of the DOPC vesicles can be estimated quite precisely with an error to about 0.2 Å. To further reduce the impact of noise in the micrograph and the limitation of pixel accuracy, we smoothened the estimated thickness values along the contour with a Gaussian kernel of sigma 2 (Fehler! Verweisquelle konnte nicht gefunden werden.**B**). The analysis of the 500 vesicles yielded a mean bilayer thickness of 28.2 Å and a standard deviation of 1.8 Å. The higher standard deviation than the estimated error above is likely due to actual variation in membrane thickness. In previous studies on DOPC bilayer thickness estimation of 50-60 non-spherical vesicles recorded at the same defocus of 2.0 µm errors were estimated between 3.0 and 4.0 Å^24^. In comparison, we obtain a slightly smaller but still very similar variance in the thickness.

### Analysis of multiple liposome test data sets

In order to comprehensively test the capabilities of CryoVIA, we compared multiple size preparations of unilamellar vesicles made of DOPC and DOPG/DPPG (70/30) lipids that are known to give rise to different ultrastructures. For the first sample, we considered the sonicated DOPC vesicles, and for the second and third, we assigned DOPC 50 nm and 200 nm size-filtered preparations, respectively. For the fourth sample, we generated vesicles from a DOPG/DPPG (70/30) mixture. For each of the samples, we prepared plunge-frozen cryo-samples and recorded between 251 and 909 micrographs (**Table 1**). Each data set was segmented by a U-net specifically trained to detect densely packed vesicles. In the four data sets, a total of 48,938 vesicles were reliably identified, while only fully closed membrane structures were considered. The detected vesicles of each data set were manually examined and discontinuously segmented or incorrectly identified vesicular structures were discarded. Incorrectly detected membranes often have unrealistic sharp bents in the contour and can, therefore, be identified by their very high curvature values. The GUI provides an easy way for sorting according to high curvature and quickly removing the segmentations by a threshold value. For each data set, the average amount of detected vesicles per micrograph varied. In micrographs of sonicated DOPC vesicles, the highest frequency of vesicles per micrograph (42.8 on average) were found due to the small circular vesicles. For the 50 and 200 nm DOPC data sets, the micrographs were often densely packed while the toolkit managed to identify on average 24.1 and 5.1 vesicles per micrograph, respectively whereas the DOPG/DPPG vesicle preparation only had an average of 4.2 vesicles per micrograph.

**Table 1.** Sample and data acquisition details of test data sets.

|  | DOPC sonicated | DOPC 50 nm size-filtered | DOPC 200 nm size-filtered | DOPG/DPPG (70/30) |
| --- | --- | --- | --- | --- |
| Lipid Concentration (mg/ml) | 5 | 5 | 5 | 5 |
| Magnification (x 10,000) | 49 | 49 | 49 | 53 |
| Pixel size (Å/pix) | 1.737 | 1.737 | 1.737 | 1.737 |
| # Frames | 45 | 45 | 45 | 45 |
| Total Dose (e<sup>-</sup>/Å<sup>2</sup>) | 45 | 45 | 45 | 45 |
| Underfocus (µm) | 2.0 | 2.0 | 2.0 | 2.0 |
| # Movies | 909 | 251 | 356 | 514 |
| # Vesicles detected/vesicle frequency per micrograph | 38,871/42.8 | 6061/24.1 | 1819/5.1 | 2179/4.2 |

When we computed global parameters of circumference, diameter, and area (**Table 2**, **Figure 5A, Figure S2A,B**), we found the that the determined circumferences of vesicles in the sonicated DOPC data set are on average smaller with a median circumference of 651 ± 189 Å median absolute deviation (m.a.d.) than in the data sets of the 50, 200 nm size-filtered DOPC and DOPG/DPPG at 1466 ± 579 Å m.a.d., 2263 ± 1356 Å m.a.d. and 2368 ± 1027 Å m.a.d., respectively. When inspecting the determined diameters of the 50 and 200 nm size-filtered DOPC vesicles, the median diameters differ from 465 ± 189 Å m.a.d. to 744 ± 458 Å m.a.d., respectively, which is in agreement with the chosen polycarbonate size filters for the vesicle preparations. The presence of vesicles with maximum diameters up to 3040 and 4374 Å larger than the size filters of 50 and 200 Å, respectively, reveals that a considerable amount of large diameter vesicles was not removed by the filter presumably due to the fluid and deformable properties of the lipid membranes. The sonicated DOPC vesicles showed a median diameter of 209 ± 58 Å m.a.d. while the DPPG-DOPG data set had a similarly high median diameter as the 200 nm DOPC vesicles with a median diameter of 737 ± 315 Å m.a.d. The obtained results including the deviations from the sonication-only data set are in agreement with previous studies as sonication produces on average smaller vesicles^25^. Together, CryoVIA is capable of automatically determining global size statistics of multiple lipid vesicle preparations from a large number of cryo-EM micrographs.

**Table 2.** Circumference, diameter, and area statistics of the four data sets.

|  | Maximum | Minimum | Mean | Median | Median absolute deviation |
| --- | --- | --- | --- | --- | --- |
| Circumference [Å] |  |  |  |  |  |
| DOPC sonicated | 15,897 | 388 | 961 | 654 | 189 |
| DOPC 50 nm size-filtered | 8,563 | 402 | 1,758 | 1,466 | 579 |
| DOPC 200 nm size-filtered | 12,566 | 402 | 3,117 | 2,263 | 1,356 |
| DOPG/DPPG (70/30) | 23,473 | 406 | 3,077 | 2,368 | 1,027 |
| Diameter [Å] |  |  |  |  |  |
| DOPC sonicated | 6,052 | 124 | 309 | 209 | 58 |
| DOPC 50 nm size-filtered | 3,040 | 126 | 575 | 465 | 189 |
| DOPC 200 nm size-filtered | 4,374 | 126 | 1,041 | 744 | 458 |
| DOPG/DPPG (70/30) | 7,364 | 134 | 960 | 737 | 315 |
| Area [nm<sup>2</sup>] |  |  |  |  |  |
| DOPC sonicated | 125,508 | 104 | 1,102 | 302 | 154 |
| DOPC 50 nm size-filtered | 41,469 | 106 | 2,875 | 1,495 | 1,027 |
| DOPC 200 nm size-filtered | 89,825 | 114 | 9,941 | 3,453 | 3,020 |
| DOPG/DPPG (70/30) | 392,323 | 118 | 10,793 | 3,995 | 2,979 |

In order to further characterize the local membrane properties of the chosen test samples, we performed detailed membrane analysis as part of CryoVIA. As a result, we found that the average of the mean bilayer thickness of sonicated DOPC vesicles and the DOPG/DPPG vesicles with 27.5 Å and 27.2 Å was smaller than in the 50 and 200 size-filtered DOPC samples at 29.0 and 30.5 Å (Fehler! Verweisquelle konnte nicht gefunden werden.**B**), which is likely a direct result of the filter treatment of sonicated DOPC vesicles. For DOPG/DPPG vesicles, the observed difference between the maximum and minimum bilayer thickness was higher at 22 Å than for the DOPC vesicles at 15 Å (Fehler! Verweisquelle konnte nicht gefunden werden.**C**), suggesting larger local structural changes, possibly due to the presence of unsaturated DPPG molecules. Inspection of DOPG/DPPG vesicles identified with high bilayer thickness supports the measured quantities visually (Fehler! Verweisquelle konnte nicht gefunden werden.**D**). Relating the bilayer thickness with diameter revealed an overall correlation between the two quantities for the vesicles taken from the four data sets (Fehler! Verweisquelle konnte nicht gefunden werden.**E**). Interestingly, sonicated DOPC as well as DOPG/DPPG samples showed a more focused distribution of diameter with bilayer thickness whereas the size-filtered data sets show a more continuous distribution of vesicles.

In order to further characterize the shapes of the four vesicular samples, we clustered them using the pre-trained default CryoVIA shape classifier. For the four samples, the most prominent shape was the spherical vesicle, in particular for sonicated DOPC and DOPG/DPPG vesicles as they made up more than 95% of all vesicles (Fehler! Verweisquelle konnte nicht gefunden werden.**A**). The 50 and 200 nm size-filtered DOPC samples showed more shape diversity as other shapes took up more than 13 and 20% share, respectively. The majority of the remaining vesicles were classified as prolates that tended to be close to spherical as well (Fehler! Verweisquelle konnte nicht gefunden werden.**B**). Examples of classified shapes were inspected and overlayed onto each other (Fehler! Verweisquelle konnte nicht gefunden werden.**C/D**) as well as a small share of unusual shapes not found in the default training set (**Figure S2C**). The complete analysis of the 2031 micrographs took 3.5 hours and a total of 48,930 vesicles were identified and analyzed (see STAR methods). In conclusion, CryoVIA reliably extracted local membrane characteristics and consistently classified the majority of the shapes of the four differently prepared vesicle data sets.

### Membrane remodeling effects of bacterial ESCRT-III member PspA

To test further the utility of the developed CryoVIA tools on cryo-EM data obtained from protein lipid mixtures, we turned to a previously characterized preparation of small unilamellar vesicles (SUVs) from *E. coli* polar lipid extract (EPL) incubated with the bacterial ESCRT-III protein PspA^8^. Previously, it was observed that upon addition of PspA the SUVs were converted into larger vesicles in comparison with the control of EPL alone and that bilayer thickness increased for large vesicles. This time, we applied CryoVIA to the two data sets, respectively (**Table 3**).

**Table 3.**
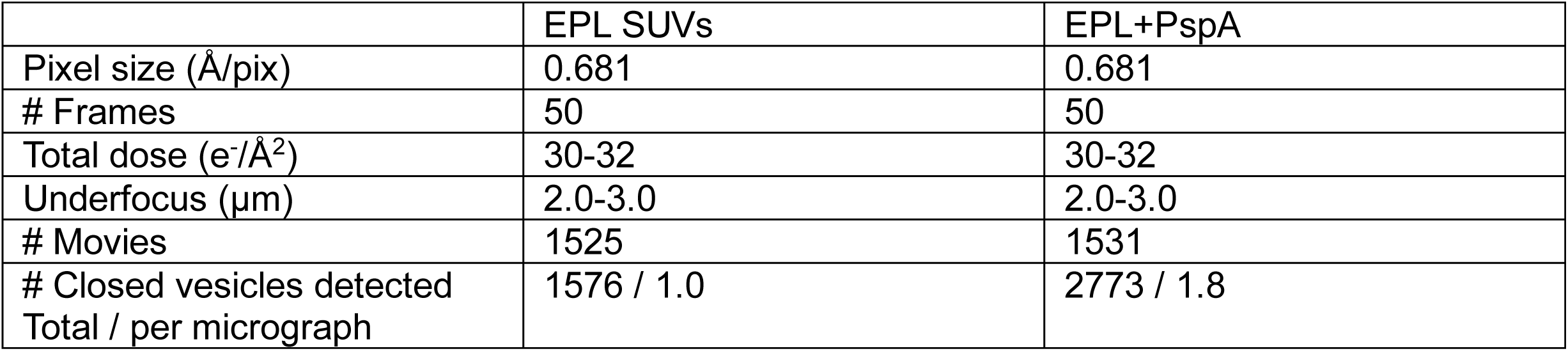
Sample and cryo-EM data acquisition details of EPL SUVs and the EPL+PspA mixture^8^.

|  | EPL SUVs | EPL+PspA |
| --- | --- | --- |
| Pixel size (Å/pix) | 0.681 | 0.681 |
| # Frames | 50 | 50 |
| Total dose (e<sup>-</sup>/Å<sup>2</sup>) | 30-32 | 30-32 |
| Underfocus (µm) | 2.0-3.0 | 2.0-3.0 |
| # Movies | 1525 | 1531 |
| # Closed vesicles detected<br>Total / per micrograph | 1576 / 1.0 | 2773 / 1.8 |

Initially, we benchmarked the capabilities of the pre-trained network. Prior to segmentation, high-pass filtering was applied to compensate for the strong contrast effects of the lacey carbon foil. This way, a total of 50 randomly selected micrographs were segmented with the provided pre-trained network. Based on the 160 manually curated reference vesicles, the pre-trained neural network detected 146 correctly while only 8 had been missing, reaching an accuracy of 0.91. The small fraction of missing vesicles is composed of non-closed vesicles. While the average circumference of membranes was similar between EPL SUVs and EPL + PspA samples, the data set containing PspA generated membranes of larger lengths (**Figure 7A**). When focusing the analysis on membranes with a circumference greater than 5000 Å (**Figure S2D**), the difference between the largest vesicles/membranes in both data sets became more apparent. While the length of EPL SUV membranes rarely exceeded 6000 Å, the EPL membranes including PspA were mostly found evenly distributed between 5000 and 8000 Å with a tail fraction even longer than 10,000 Å. This analysis was limited by the fact that a lot of the membrane structures were larger than the dimensions of the micrographs and were, therefore, not automatically analyzed due to the non-closed entity present on the micrograph (**Figure S3A**). By contrast, in the respective publication^8^, the authors had annotated membranes with a length of over 15,000 Å by fitting an ellipsoid shaped vesicles the beyond the micrograph, which we did not implement in CryoVIA. Nevertheless, the clear trend in size difference between EPL SUVs and EPL+PspA as a result of PspA-mediated membrane fusion was confirmed by the CryoVIA workflow.

When analyzing the mean bilayer thickness of the two data sets (Fehler! Verweisquelle konnte nicht gefunden werden.**B**), at first sight the bulk distribution looked very similar with a mean thickness of around 30 Å and most values found 25-35 Å. Nevertheless, the EPL+PspA sample had an additional distribution of membranes with a bilayer thickness of around 45-55 Å. When we examined the bilayer thickness of membranes with a length greater than 5000 Å (**Figure S2D/E**), we detected a shift towards a greater thickness in the PspA data set with a mean of 42 Å compared to the new mean of 33 Å in the EPL SUV data set, thus confirming the reported conclusions that larger vesicles on average possess an increased bilayer thickness^8^. Moreover, when correlating the mean bilayer thickness with circumference a second population emerged with bilayer thickness greater than about 45 Å mostly found in vesicles of large circumference (Fehler! Verweisquelle konnte nicht gefunden werden.**C/D**). It should be noted most of the large circumference membranes with a thicker bilayer exceeded the dimensions the micrograph (**Figure S3B**) and thus limiting the precise reporting of this apparent structural property. For more precise reporting and analysis through CryoVIA, lower magnification micrographs with larger fields of views will more accurately capture structural changes of large membrane entities. We demonstrated that CryoVIA can be successfully employed to protein lipid mixtures in order to characterize the protein-induced membrane remodeling effects on lipids.

## Discussion

We introduced a dedicated Cryo Vesicle Image Analyser (CryoVIA) software package for the evaluation of membrane structures in cryo-EM data sets. CryoVIA comprises the complete image analysis workflow from neural-network segmentation, membrane structure identification followed by the structure analysis (**Fehler! Verweisquelle konnte nicht gefunden werden.**). When complete membrane structures are identified, they are subjected to the analysis of local and global structural parameters such as membrane thickness and curvature with high precision as well as diameter, circumference, and area followed by classification into distinct membrane shapes (**Figures 2, 3, 4**). CryoVIA was tested on the analysis of four different vesicle data sets confirming relevant differences induced by the preparation procedure and membrane composition (**Tables 1, 2**; **Figures 5, 6**). Moreover, CryoVIA was applied to quantitively evaluate membrane shape changes of a previously reported bacterial ESCRT-III membrane remodeling protein PspA (**Table 3, Fehler! Verweisquelle konnte nicht gefunden werden.**).

**Figure 1.**
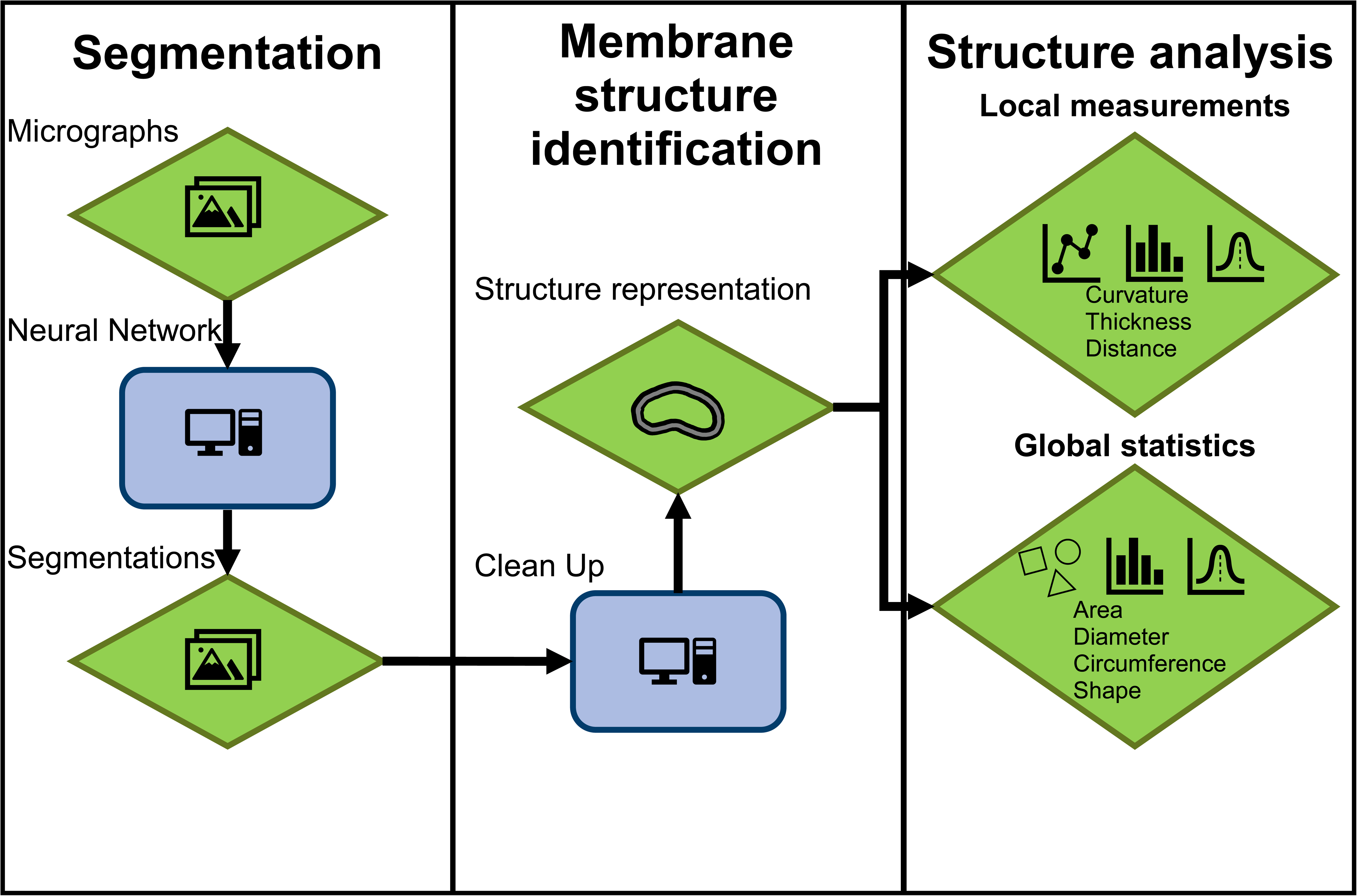
Workflow of the cryo vesicle image analyzer (CryoVIA). Cryo-micrographs serve as input for the U-net based segmentation that is followed by a membrane structure identification. This step assigns unique and single membrane entities and is critical to single out overlapping membrane structures. In the final step, the identified membrane structures are subjected to a detailed quantitative analysis for the extraction of local and global structural parameters.

**Figure 2.**
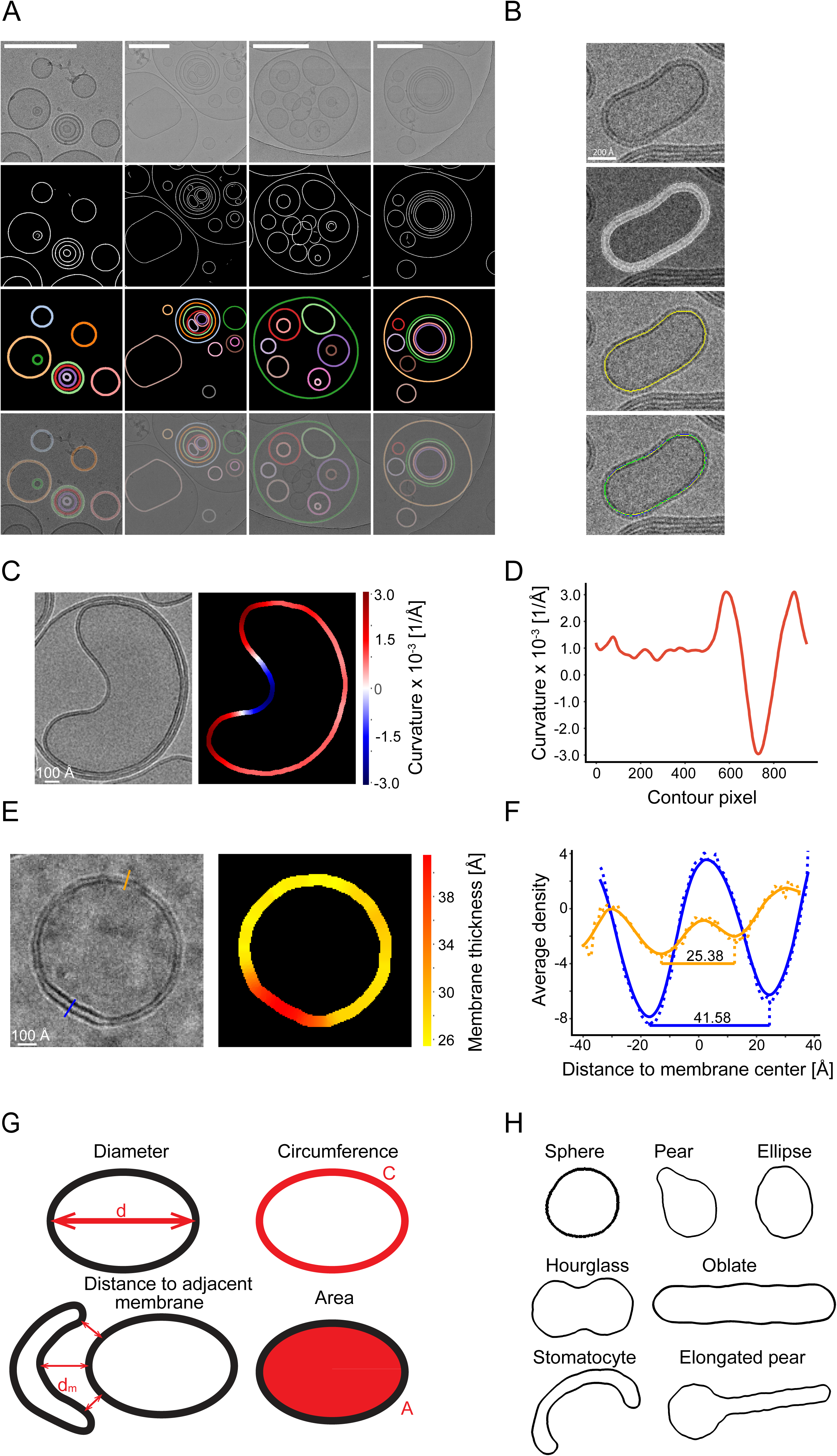
CryoVIA’s main functions: membrane segmentation followed by structural parameter and shape analysis. (A) From top to bottom rows: Four micrographs of the data set DOPC (200 nm size filtered) containing different lipid structures (top). The corresponding U-net based segmentation (top center). The results of membrane identification (bottom center). Segmentation results superimposed on original micrographs (bottom). Only closed vesicles were extracted from the segmentation. Scale bar 200 nm. (B) From top to bottom: Magnified image of vesicle (top). Segmentation of the corresponding vesicle: multiple pixels are covering the bilayer (top center). One pixel-wide initial skeleton of the segmentation in yellow (top bottom). Skeleton in yellow, corrected skeleton in blue, pixels where skeleton and corrected skeletons are identical in green (bottom). (C) Example stomatocyte vesicle from the DOPC 200 nm size-filtered data set (left) with curvature values superimposed on segmented membrane (right). (D) Curvature of C plotted as a function of contour pixels. (E) Example vesicle from the DOPG/DPPG data set with two cross sections in blue and orange (left) with membrane thickness estimates superimposed on segmented membrane (right). (F) Extracted (dashed line) and smoothened (solid) density profiles from the corresponding cross section locations in E blue and orange, respectively. (G) Global parameters extracted are diameter, circumference, distance to adjacent membranes, and area. (H) Possible membrane shapes from membrane structures are: spheres, pears, ellipsoid, hourglass, oblate, stomatocyte, and elongated pear. These shapes are used for the training of the default shape classifier later (see also Figure S1 and S2).

**Figure 3.**
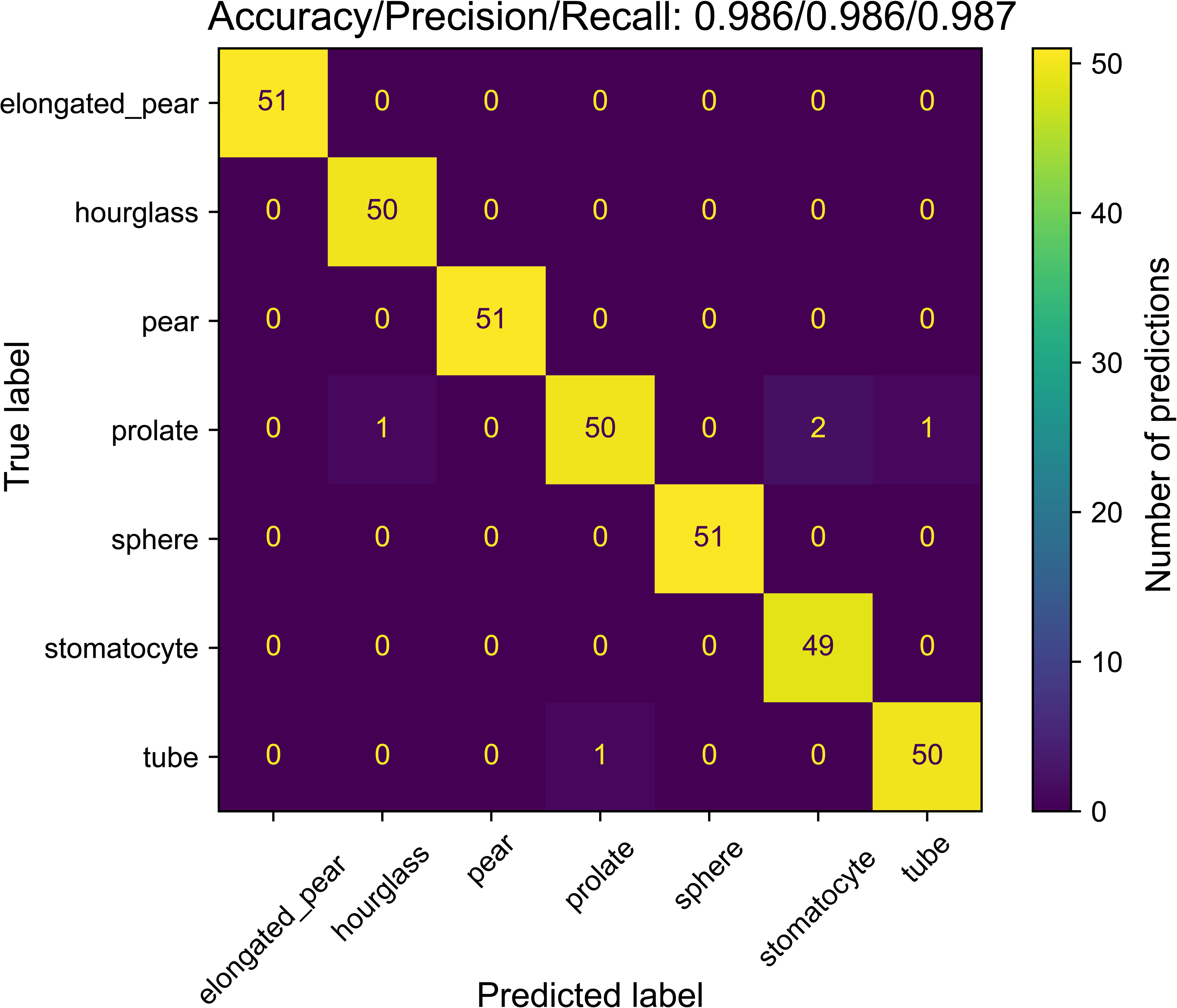
Performance indicators of shape classification. Confusion matrix and classification metrics accuracy, precision, and recall for the default shape classifier trained on seven different shapes.

**Figure 4.**
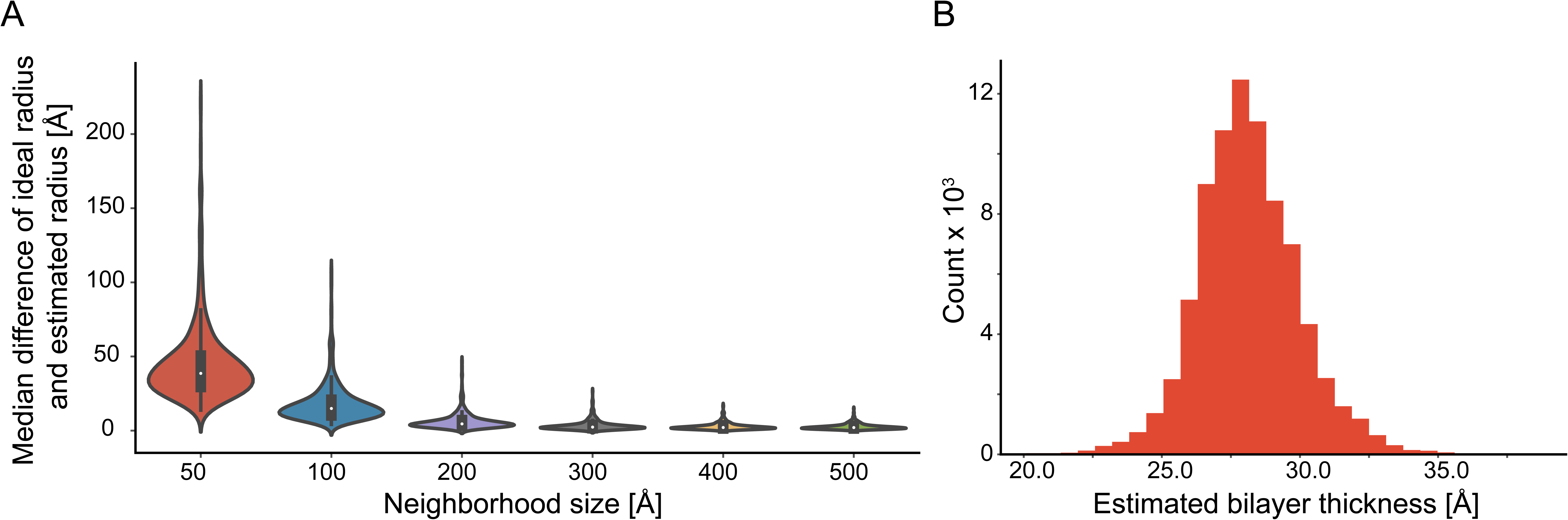
Error estimation of curvature and bilayer thickness determination in a DOPC sonicated data set using the 500 most circular vesicle contours found. (A) Median of difference between the globally estimated vesicle radius (ideal) and the locally determined (estimated) curvature radius using different maximum neighborhood sizes for curvature estimation. For neighborhood sizes larger than 200 Å the errors are smaller than 6 Å. (B) Bilayer thickness estimations of all contour points yield a distribution of 28.2 ± 1.8 Å. One standard deviation of the mean accounts for 72.4 % of included values.

**Figure 5.**
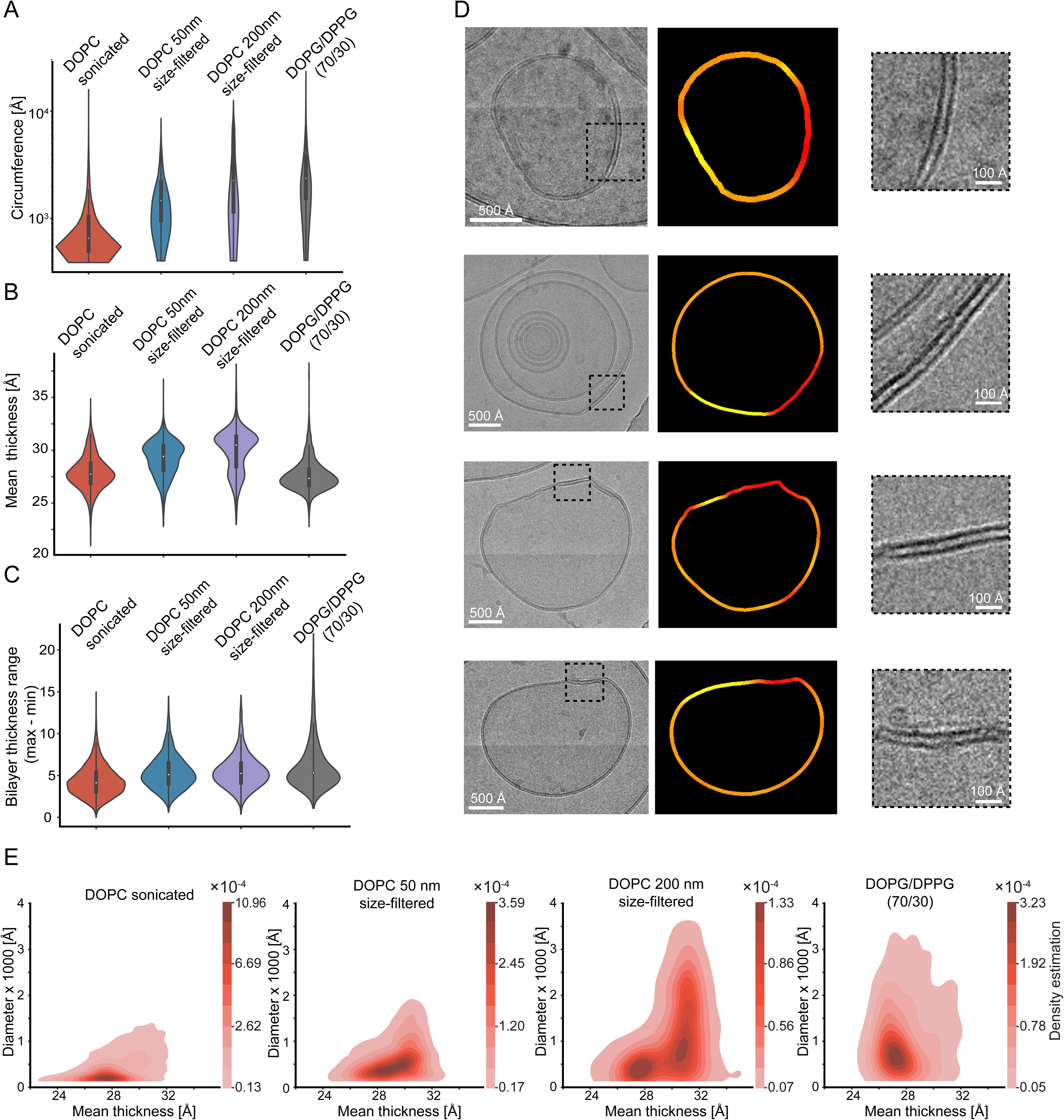
Determined quantities of circumference, diameters, and membrane thickness from four lipid samples. (A - C) Violin plots of the evaluated vesicles from four data sets (DOPC sonicated (n = 38871), DOPC 50 nm size-filtered (n = 6061), DOPC 200 nm size-filtered (n = 1819), and DOPG/DPPG (70/30) (n = 2179)) with circumference (A), mean bilayer thickness distribution and range (C). The white dot within each violin indicates the median, while the shaded area represents the kernel density estimate of the data distribution. The inner grey box represents the 25% to 75% interquartile range, while the vertical grey line includes values inside the interquartile range multiplied by 1.5. (D) Example images for high bilayer thickness of the DOPG/DPPG data set. The vesicle image with a dashed box indicating the region of zoomed inset (left). The thickness map along the segmented contour from yellow to red with red being the highest thickness (center). Zoomed membrane segment with the highest thickness value (right). (E) Kernel density distribution plots of diameter against the mean bilayer thickness. The density values of each distribution are normalized in such a way that the sum equals 1 (see also Figure S2).

**Figure 6.**
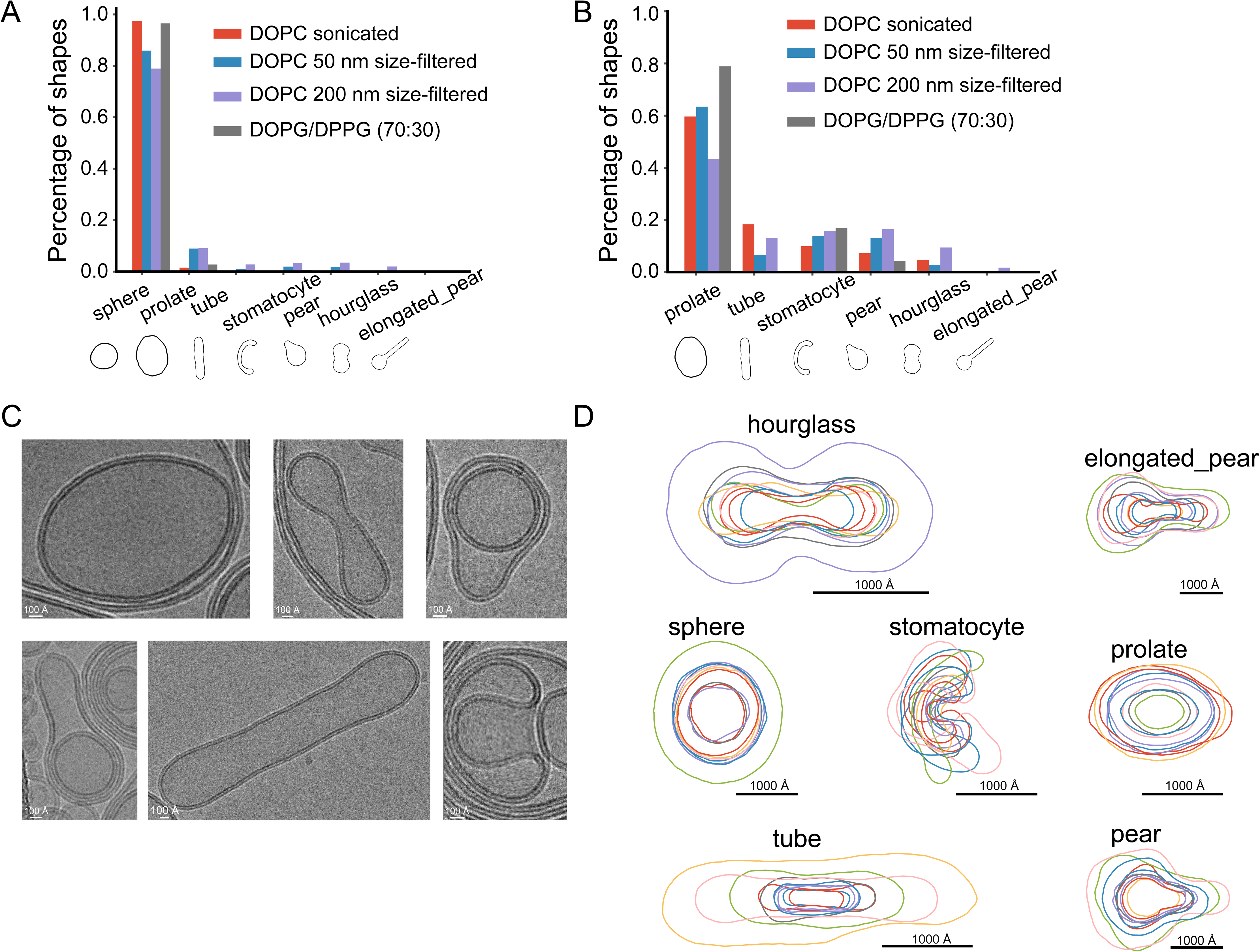
Shape classification of sonicated, 50 and 200 nm size-filtered DOPC and DOPG/DPPG vesicles. (A) Bar graph of the shape distribution of the four data sets. (B) Bar graph of the shape distribution of all data sets excluding spheres as they make up the majority of shapes. (C) Example vesicle images of shapes found in micrographs. From top left to bottom right: prolate, hourglass, pear (from DOPC 200 nm sample, respectively), elongated pear (DOPC sonicated), tube (DOPC 200 nm) and stomatocyte (DOPC sonicated). (D) A total of 7 classified shapes, 10 aligned and superimposed experimentally segmented vesicle contours. The 10 vesicles with the highest confidence for each shape in the shape classification are displayed (see also Figure S2).

**Figure 7.**
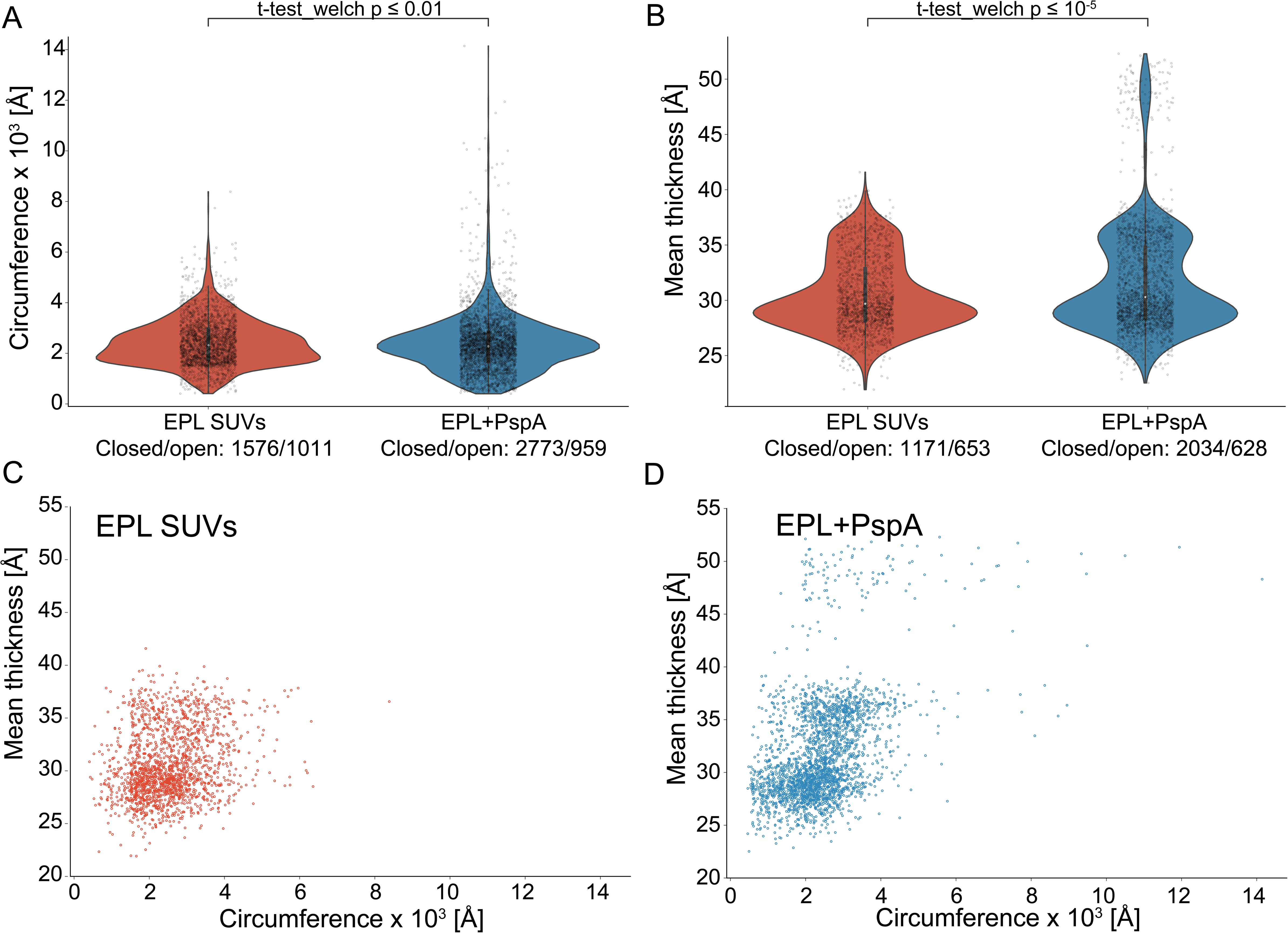
Membrane remodeling analysis of the EPL+PspA data set. (A) Violin plot of the membrane length distribution. (B) Violin plot of the mean bilayer thickness distribution. (A, B) The white dot within each violin indicates the median, while the shaded area represents the kernel density estimate of the data distribution. The inner grey box represents the 25% to 75% interquartile range, while the vertical grey line includes values inside the interquartile range multiplied by 1.5. The sample size is noted below the label. (C) Scatter plot of mean bilayer thickness against circumference of the EPL SUVs data set (D) Scatter plot of mean bilayer thickness against circumference of the EPL + PspA data set (see also Figure S3).

The here obtained membrane thicknesses were determined directly from the cryo-EM images with an estimated spatial precision of ± 0.2 Å. When image intensities are used for thickness estimation, they interestingly correspond to the lipid hydrophobic thickness (2D_C_)^17^. This thickness is given by the distance covering the lipid chains embedded present in the bilayer and is slightly lower than the expected distance between the denser phosphate head groups (D_HH_) that are commonly obtained by X-ray or neutron scattering methods. For instance, in the case of DOPC (50 nm size filtered) the 2D_C_ and D_HH_ were determined by SAXS at 29.3 and 35.0 Å, respectively. The same study reported cryo-EM based measured bilayer thickness of DOPC lipids (50 nm size filtered): mean thickness value of 29.7 ± 0.6 Å matching the hydrophobic thickness (2D_C_)^17^. The here estimated bilayer thickness of the DOPC 50 nm size-filtered test dataset at 29.0 Å compares well with the reported cryo-EM value of 29.7 Å. The slight differences are likely explained by different pixel size calibrations. Reasons for the deviation to the actual phosphate head group distance D_HH_ arise from the smearing of the obtained lipid intensities due to the 2D projection through a spherical 3D lipid membrane convolved with the CTF. For DOPG and DPPG, the lipid hydrophobic thicknesses 2D_C_ were reported to be 27.2 Å and 26.8 Å, respectively, albeit at elevated temperatures of 50°C^26^. For the mean bilayer thickness of DOPG/DPPG using the CryoVIA method, we obtained 27.2 Å that is in close correspondence to the previously reported values. Together, we find the CryoVIA estimates on membrane thickness very precise on sub-Å-scale and accurate on an Å level as they are in good agreement with previously reported values by scattering methods. Regardless of minor deviations, CryoVIA can be used to reliably estimate membrane thickness in particular for comparative studies that investigate membrane thickness changes.

Imaging of vesicular lipid structures using cryo-EM has been a direct way to visualize and characterize the structures at high resolution. In some contexts, negative staining electron microscopy has been used. However, as the technique requires specimen drying followed by a collapse of the lipid structures, the detailed structural properties are not faithfully maintained for imaging. In contrast, the main advantage of cryo-EM is the structural preservation of the specimen in a fully hydrated manner after plunge-freezing them in vitreous ice. However, it should be mentioned that during the freezing procedure, the volume of the starting droplet is reduced drastically into water films of 40 - 80 nm of thickness imposing stress and crowding of the large lipid structures. This preparation procedure excludes lipid structures from the analysis that are larger than the dimensions of the ice film thickness such as giant unilamellar vesicles. Nevertheless, although the observed lipid structures of appropriate sizes may suffer from some deformation, the cryo-EM method presents a direct and faithful way of imaging lipid structures in aqueous solution.

The presented CryoVIA was developed and tested using cryo-EM micrographs of purified lipid-water mixtures. In this case, the structural analyses are restricted to the images in two dimensions. The analyzed micrographs were recorded with a dose of 20 – 30 e^-^/Å^2^ at an underfocus range between 2.0 – 3.0 µm. Previous analyses revealed that defocus series did not have an effect on the estimated bilayer thickness measurements while images close to focus suffered from poor overall contrast problematic for image analysis^24^. The precision and easy accessibility to quantifications of lipid shape changes opens up new possibilities to study membrane-remodeling proteins^8–10^ such as dynamin and ESCRT-III proteins.

Vesicle identification in cryo-micrographs can also be used to aid in solving the structure of membrane proteins using single particle structure determination^27^. Although not implemented, CryoVIA applications could also be extended to inform particle picking for single-particle structure determination of bilayer-embedded protein complexes, e.g., by selecting protein complexes that reside in vesicles of particular diameter ranges and lipid curvatures in analogy to the previously published tool Vesicle Picker^28^. Various methods for solving structures of membrane-bound proteins in liposomes have been proposed^29^. This way, proteins can be directly isolated in cell-derived membrane vesicles^30^ and exposed to additional stimuli, e.g., electric fields^31^ and buffer gradients^32,33^. Although CryoVIA was not designed for single particle protein structure determination, it could be used to enhance the homogeneity of the structure by classifying picked particles through membrane curvature or vesicle shapes. Nevertheless, the current implementation of CryoVIA focuses on analyzing the membrane and vesicle structures themselves rather than solving the structure of membrane proteins. Lipid membrane analysis tools in three dimensions have been developed for electron tomography^18,20^. We here show that many of the conclusions on ultrastructural lipid changes can already be observed in two-dimensional images at better signal-to-noise ratios without the need of tilt-series acquisition and tomographic reconstructions. Moreover, the application of zero-tilt imaging has already successfully been extended to the *in situ* cellular environment^34^ and, therefore, the analysis of lipid structures from cellular images based on two-dimensional micrographs are expected to be possible. The results of the quantitative lipid structure analysis will largely depend on the initial results of segmentation, and, therefore alternative and more tailored membrane segmentation approaches^19,35^ could be combined with the analytical tools from CryoVIA. CryoVIA’s ability to process multiple data sets efficiently streamlines the analysis process. This feature is particularly advantageous when dealing with large-scale comparative studies including high-throughput experiments.

**CryoVIA is a python-based program that is designed to run quickly on a desktop computer to run the analysis in a user-friendly manner, i.e., a data set of 2031 micrographs can be fully analyzed within four hours of runtime on a dedicated work station (see** STAR methods). One of the key advantages of our toolkit is the ability to accurately measure key membrane parameters such as bilayer thickness, diameter, and shape of vesicles. This level of precision is critical for understanding the structural and molecular variations of biological membranes of vesicles and organelles. The toolkit’s user-friendly interface and algorithms ensure that researchers, regardless of their expertise, can access and utilize these analytical tools. The presented toolkit offers a robust and accessible solution for the analysis of membranes in cryo-EM micrographs opening up new avenues to study protein lipid interactions. Moreover, the software can also be employed to address experimental challenges in adjacent fields of structural biology such as nanoparticles used in drug delivery as well as soft-matter science.

## Resource Availability

### Lead Contact

Requests for further information and resources should be directed to and will be fulfilled by the lead contact, Carsten Sachse.

### Materials Availability

This study did not generate new unique reagents.

### Data and Code Availability

All data reported in this paper will be shared by the lead contact upon request. CryoVIA is publicly available at https://github.com/philipp-schoennenbeck/CryoVia and https://doi.org/10.5281/zenodo.14335953. A GUI is implemented for ease of usage and knowledge of programming is not needed to use this software.

The four data sets of DOPC (sonicated), DOPC (50 nm size-filtered), DOPC (200 nm size-filtered) and DOPG/DPPG (70/30) have been deposited at https://www.ebi.ac.uk/empiar/ with the following access codes: EMPIAR-12520, EMPIAR-12523, EMPIAR-12521 and EMPIAR-12522, respectively. They are publicly available as of the date of publication. Accession codes are also listed in the key resources table. Any additional information required to reanalyze the data reported in this paper is available from the lead contact upon request.

## Acknowledgements

The authors thank Sabrina Berkamp for critically reading the manuscript. This study was funded by the Deutsche Forschungsgemeinschaft (DFG, German Research Foundation, SA 1882/6-1 (CS) and CRC1208 Project Nr 267205415 (CS)). The authors gratefully acknowledge the electron microscopy access time and computing time granted by the biological EM facility of the Ernst-Ruska Centre at Forschungszentrum Jülich. In this regard, we thank Thomas Heidler, Saba Shazad, and Pia Sundermeyer for maintaining the electron microscopes and Daniel Mann for maintaining the processing computers. The authors gratefully acknowledge the computing time granted by the JARA Vergabegremium and provided on the JARA Partition part of the supercomputer JURECA at Forschungszentrum Jülich^36^.

## Author contributions

P.S. and C.S. designed the research. B.J. prepared liposome samples and imaged them by the cryo-EM. P.S. wrote code for CryoVIA. P.S. and C.S. evaluated the presented data. P.S., and C.S. prepared the manuscript with input from B.J.

## Declaration of interests

The authors declare no competing interests.

## STAR methods

### Key resource table

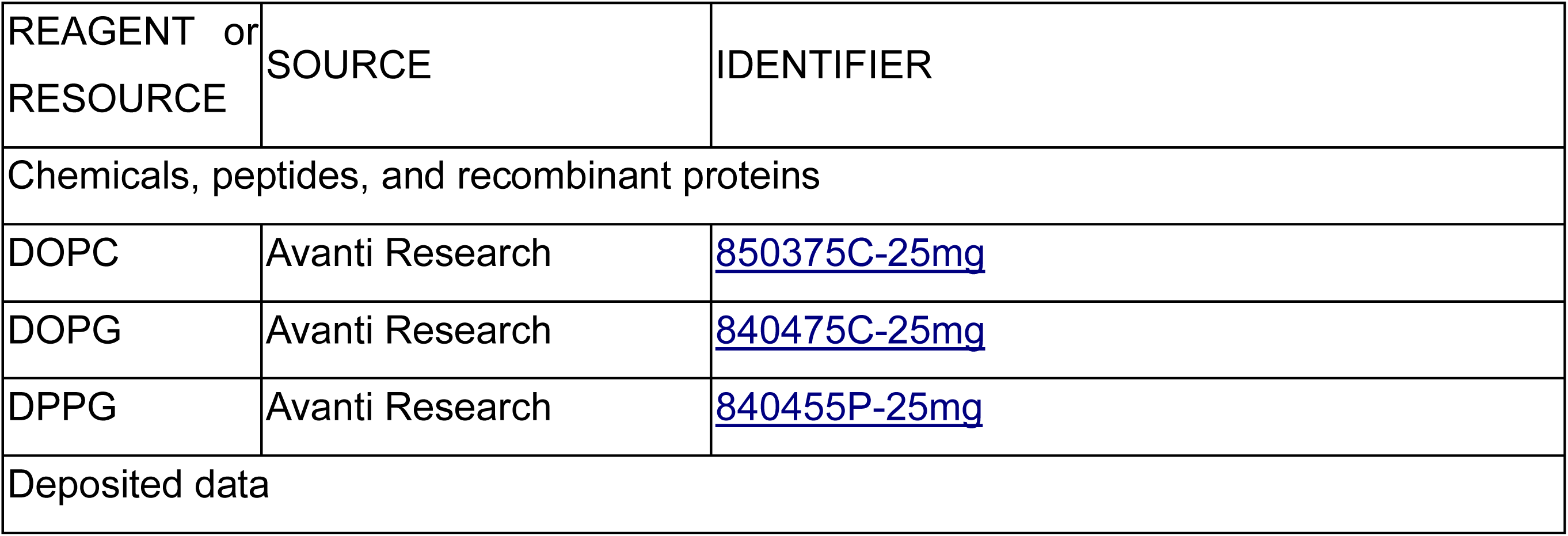
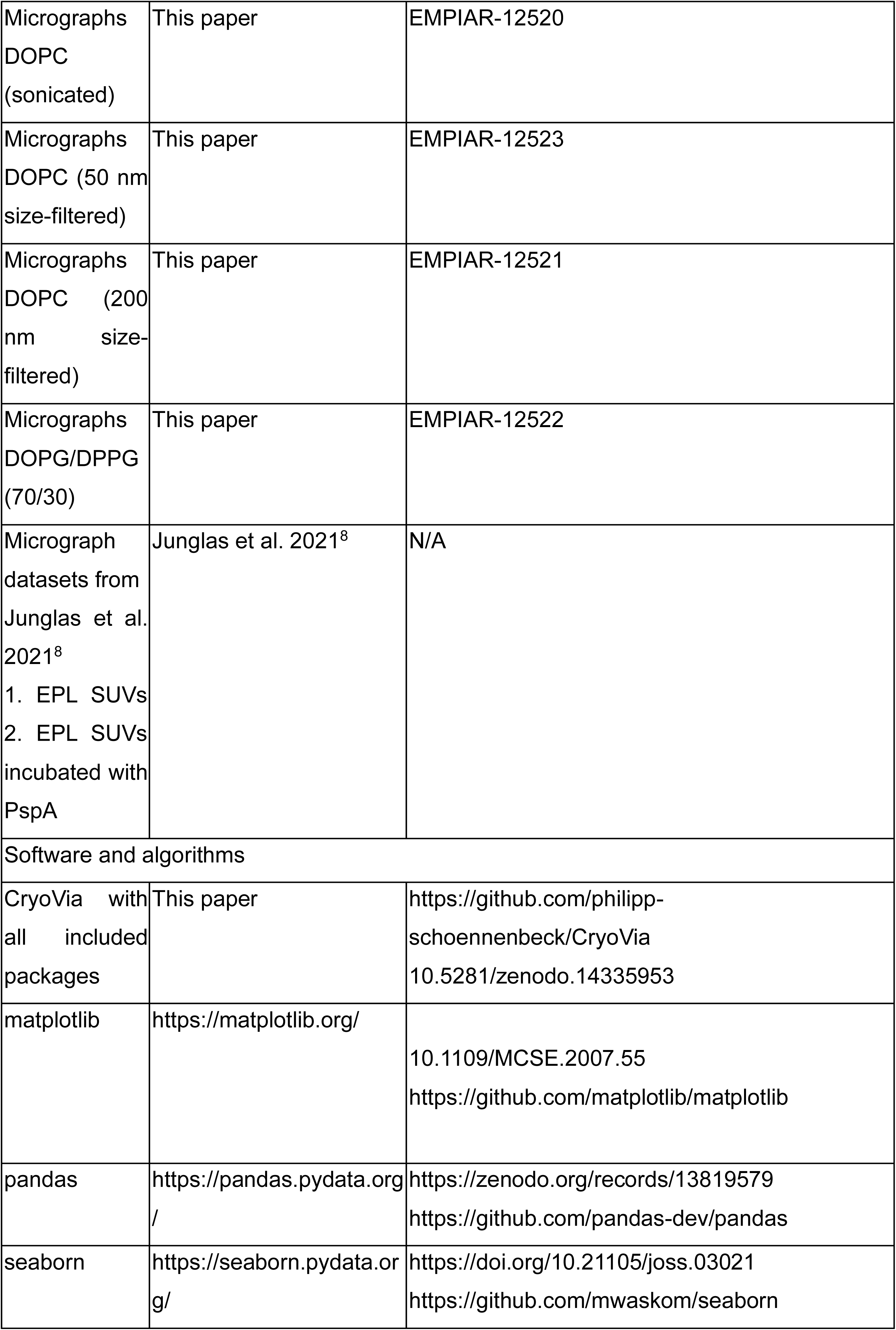
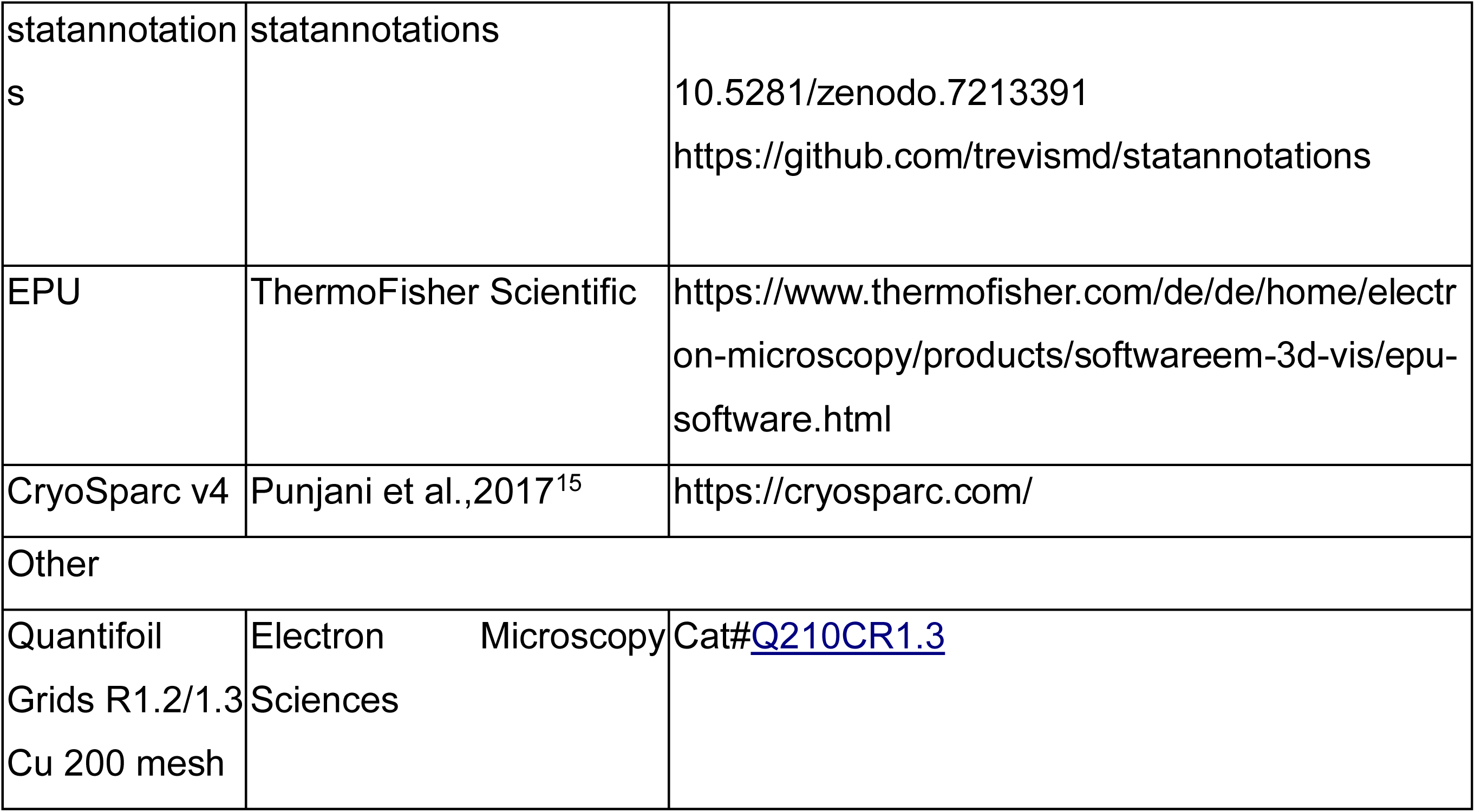

## METHOD DETAILS

### Liposome preparation

Chloroform dissolved 1,2-dioleoyl-sn-glycero-3-phosphocholine (DOPC), 1,2-dioleoyl-sn-glycero-3-phospho-(1’-rac-glycerol) (DOPG) and 1,2-dipalmitoyl-sn-glycero-3-phospho-(1’-rac-glycerol) (DPPG) were purchased from Avanti polar lipids. Lipid films were produced by mixing the lipid solutions (100% DOPC or 70% DOPG/30 % DPPG (w/w) mixture) and evaporating the solvent under a gentle stream of nitrogen and followed by vacuum desiccation overnight. The lipid films were rehydrated in 10 mM Na-Phosphate buffer pH 7.6 (DOPC) or 10 mM Tris-HCl pH 8.0 (DOPG/DPPG) by shaking for 30 mins at 37 °C (DOPC) or 50 °C (DOPG/DPPG). The resulting liposome solution was subjected to five freeze-thaw cycles. The liposome solution was either used as is (DOPG/DPPG), or SUVs (small unilamellar vesicles) were generated by extrusion of the liposome solution through a porous polycarbonate filter (50 nm pores, 200 nm pores) or tip sonication (40% power output, 50% duty cycle, 5 min) using an Ultrasonic Homogenizer (Biologics, Inc.).

### Electron cryo-microscopy

Liposome grids were prepared by applying 4 μL sample (for sample details see **Table 1**) to glow-discharged (PELCO easiGlow Glow Discharger, Ted Pella Inc.) Quantifoil grids (R1.2/1.3 Cu 200 mesh, Electron Microscopy Sciences). The grids were plunge-frozen in liquid ethane using a Leica EM GP2 set to 80% humidity at 10 °C (sensor aided backside blotting, blotting time 3 – 5 s). Movies were recorded in underfocus on a 200 kV Talos Arctica G2 (ThermoFisher Scientific) or a 300 kV Titan Krios G4 (ThermoFisher Scientific) electron microscope equipped with a Bioquantum K3 (Gatan) or Biocontinuum K3 (Gatan) detector operated by EPU (ThermoFisher Scientific). Movie frames were gain corrected, dose weighted, aligned, binned to the physical pixel size and the CTF of the micrographs was estimated using cryoSPARC Live^15^. Micrographs with poor ice or poor CTF fit were removed.

### Segmentation

Segmentation was performed by a U-Net with image patches of 256 x 256 pixels, a depth of 4, 32 filters in the first layer, a kernel size of 3, 2 convolution layers per depth and a training rate of 4*10^-4^. A total of 92 images were manually segmented and used as training data with a pixel size of 7 Å per pixel. As the micrographs of the analyzed data sets contained a large number of overlapping and very closely packed membranes, the following specific strategy was applied. Manually segmented images were passed to the contour improvement algorithm and only corrected contours were used for the training of the U-Net. To obtain a consistent thin segmentation centered at the lipid bilayer, the contour pixels were extracted and dilated for one iteration. For images of less densely packed membranes, this step is not required and not advisable as it removes a lot of the training data. Four different segmentation strategies were tested and evaluated for their recall rates (Fehler! Verweisquelle konnte nicht gefunden werden.). While the overall metrics ranked strategy I using all images highest, we still decided for strategy IV as the results were robust for separating densely packed membranes in micrographs and provided thinner segmentation contours.

As part of feature identification, the segmented tracks need to be pruned in particular to disentangle overlapping membrane structures. For this step, the connected components of the segmentation were extracted and for each component the skeleton was calculated (**Figure S4A**). Subsequently, the skeleton was converted to a graph network. According to graph theory, nodes represent the crossovers of membrane skeletons and edges represent the membrane skeleton segments connected to crossover nodes. (**Figure S4B/C**). Each node was then evaluated to connect membrane segments that may be part of the same membrane by calculating the angle between the segments. When this angle was larger than 125°, the segments were not combined whereas when two segments formed a straight line, the skeletons were merged into one continuous membrane structure. The segment combination that did not form a straight line but had an angle smaller than 125° was evaluated by the curvature and the corresponding center of the fitted circles (**Figure S4D/E**).

The segments were combined when these values were similar. The output of this method is a binary segmentation mask for each individual membrane (**Figure S4F**). In this approach, overlapping membrane segments can be assigned to continuous underlying membrane stretches.

For subsequent parameter determination, the refinement of the membrane bilayer center is critical as the segmentation procedure inconsistently detected inner and outer leaflet separately. In order to remove those ambiguities, we convolved the micrograph with a bilayer-like kernel consisting of a ring where the outer radius represents half the maximum bilayer thickness and the inner radius was chosen such that the distance between the outer and inner radius corresponded to the leaflet thickness. As a result of this convolution, the bilayer membrane signal was enhanced and presented by a single line that was boosted by the application of the Frangi filter, which had been developed to enhance vessel-like structures^22^. Lastly, the detected lines were further simplified using interpolation through selected points of highest pixel intensities of the filtered image. When this method of detecting the bilayer center failed due to weak signal, the segmented membranes were labeled unsuitable for more detailed membrane bilayer thickness analysis. In order to detect grid holes for feature removal, the following steps were preformed: given the priorly known hole size of, e.g., 1.2 or 2.0 µm, we created a binary reference circle with the appropriate size of the grid hole. Subsequently, we performed a convolution of the micrograph with the reference image and calculate the difference between the mean pixel values outside of the circle and inside of the circle for every possible circle coordinate. The coordinate with the highest difference was the most likely spot for the center of the grid hole.

### Curvature Estimation

The local curvature is calculated at each contour point of the membranes. For each point *j*, a neighborhood *M_j_* is extracted:

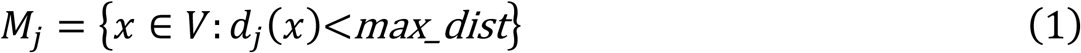

where *V* is the set of all points assigned to the current membrane, *d_j_(x)* is the distance between *j* and *x* along the contour of the membrane, and *max_dist* is a set variable that affects the sensitivity of the curvature estimation. For closed membranes, the maximum value for *max_dist* is 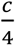, where *c* is the circumference of the closed membrane.

In the next step, the curvature *curv_j_* at each point *j* is estimated by

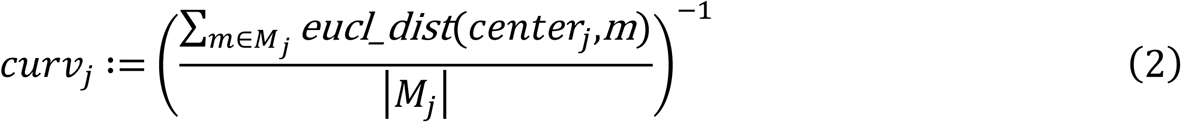

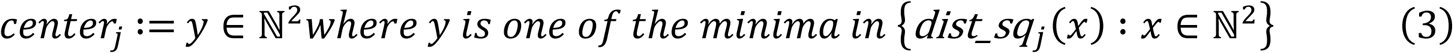

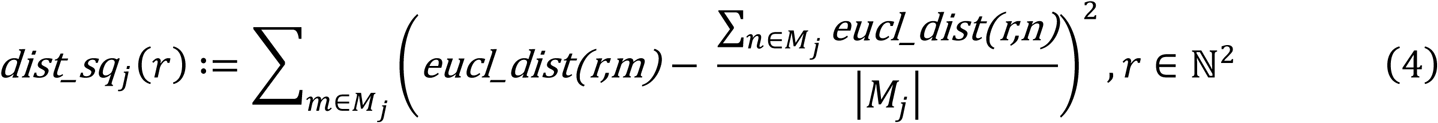

where *eucl_dist(x, y)* is the Euclidean distance between *x* and *y*, *center_j_* is the center of the best-fitting circle for the neighborhood of point *j*, and *dist_sq(r)* is the sum of squares of the differences between the Euclidean distances of *r* and *x*, and the mean of the Euclidean distances between all points of *M_j_* and *r* for all points *x* in *M_j_*. The calculation of the sign of the curvature can only be reliably determined when the membrane is part of a closed structure. For this scenario, the calculation was performed for each point *j* by taking a small step in the direction of the *center_j_*. When the resulting point was inside the closed vesicle, the curvature was considered positive and when the resulting point is outside the closed membrane, the curvature is considered negative. For non-closed membranes, determining the sign of the curvature value was estimated such that the vector from *j* towards the *center_j_* was on the same side of the tangent vector as the direction towards the previously estimated normal vector. An illustration of the curvature estimation is shown in **Figure S4G**.

### Shape classification

Vesicles were found in varying sizes and orientations on the micrographs. As the curvature decreases with increasing vesicle size, vesicles must be normalized to have consistent values for a one-dimensional convolutional neural network classifier. As a first step, the vesicles were rotated such that the two most distant points of the contour was aligned horizontally by rotating the vesicle. After rotation, the vesicle was also resized to a fixed size of 200 pixels between two points used for rotation. The ratio of the new vesicle width to the originally rotated vesicle width was then used to normalize previously calculated curvature values. To provide a consistent input to the classifier, the curvature contour was interpolated to 200 values. For the classification, the contour was shifted to the smallest curvature value at the beginning and reversed when the first maximum appeared in opposite direction.

### Bilayer thickness estimation

As the segmentation and the contour detection are usually performed on a lower resolution micrograph (the default network segments at 7 Å/pixel), the contour has to be refined to the original resolution of the micrograph. A modified procedure based on local pixel averaging from Heberle et al.^17^ was employed: After interpolating the contour pixels onto the original micrograph size, the distance between the contour pixel and all pixels surrounding the contour pixels was estimated up to a predefined maximum distance. The distance of pixels within the vesicle was considered as negative to differentiate between the different directions. In the next step, all pixel density values for the same distance bin were smoothened with a Gaussian filter with a sigma value of 10 Å, averaged and plotted as a function of distance from the contour. The values were smoothened by a Gaussian filter until two distinct minima can be extracted on either side of the contour and their distance taken as the bilayer thickness.

### Graphical user interface

CryoVIA is implemented in python and is available as a graphical user interface (GUI) for ease of use. The GUI is split into four sections:

1. Segmentation training (**Figure S5A**) In the first section, the user can create, copy or delete specific neural networks models. The default model is trained on the data sets we analyzed but additional models can be created by training a completely blank model or fine-tuning existing models. Depending on the variance in different data sets, it is helpful to use different models for different data sets. The segmentation section also provides a method to manually annotate training data by opening a plugin in napari^37^.
2. Shape classifier training (**Figure S5B**) The second section can be used to create new shape classifier models. The user can add and remove specific shapes from models and create new shapes by drawing in the provided window.
3. Foil hole Edge detection (**Figure S6A**) The third section can be used to detect and exclude carbon outside of foil holes in micrographs. This detection can be directly applied to already created data sets or the correct parameters can be identified and applied during the data set analysis. When detection is applied to data sets, membranes found outside of the foil holes will be removed from the data sets. This task is also available as a standalone software and can be applied to particle picking jobs of Relion^38^ or CryoSparc^15^.
4. Membrane analysis (**Figure S6B**) The last section is the main part of the GUI and can be used to create new data sets, run segmentation and analysis on data sets and to inspect and clean data sets. The inspection offers various filters and possibilities to inspect single membranes found in the data sets. It is also possible to compare the results of different data sets and export various results as csv-files.

### Computational resources

For the analysis of the four datasets, we used five threads with 10 parallel jobs on a server with 56 Intel(R) Xeon(R) Gold 5120 CPU @ 2.20GHz processors and two Nvidia Geforce RTX 4080 GPUs for the segmentation. The segmentation including the structure identification and the analysis of the four datasets with a total of 2031 micrographs took 3.5 hours yielding a total of 50,561 vesicles. The segmentation and the analysis took 72 and 138 minutes, respectively. On average, the segmentation of a micrograph took 2.1 s and the analysis of a single vesicle 0.2 s. These values varied depending on the hardware, the size of the micrographs, the number of vesicles found per micrograph and the size of the vesicles. The 50,561 identified vesicles were reduced to 48,930 by using filtering and manual clean up provided by the GUI.

## QUANTIFICATION AND STATISTICAL ANALYSIS

The relevant data columns were exported from CryoVIA and figures were created using matplotlib^39^, seaborn^40^, pandas^41^, statsannoations^42^. To evaluate the accuracy of the default vesicle shape classifier we used the standard definition of accuracy, precision and recall. The accuracy of the curvature calculation with different neighborhood sizes was estimated by calculating the difference between the estimated radii and the ideal radii for the most circular segmented vesicles and visualize the median for each of these vesicles. The same vesicles were used to visualize the distribution of the estimated bilayer thickness to show a high correlation to a Gaussian distribution. Correlation between the vesicle diameter and bilayer thickness was visualized by a kernel density estimation plot using the python seaborn package. To show differences between the circumference and mean thickness distribution of the EPL SUVs and EPL+PspA datasets a Welch’s t-test was calculated by the python package statannotations.

## Notes

https://github.com/philipp-schoennenbeck/CryoVia

### Competing Interest Statement

The authors have declared no competing interest.

### Summary of Updates

Fixed some errors in the citations.

