## Supplemental Figures and Tables for "CryoVIA - An image analysis toolkit for the quantification of membrane structures from cryo-EM micrographs"

**List of supplemental items**

**Table S1. Recall rates of four segmentation strategies for 11 random test micrographs, Related to STAR Methods.**

| | I<br>All images | II<br>All images + thinning | III<br>Only contour improved membranes | IV<br>Only contour improved membranes + thinning |
|---|---|---|---|---|
| Accuracy | 0.96 | 0.97 | 0.97 | 0.99 |
| Mean Precision | 0.85 | 0.73 | 0.85 | 0.79 |
| Mean Recall | 0.95 | 0.92 | 0.94 | 0.89 |
| Mean Intersection over Union | 0.81 | 0.71 | 0.81 | 0.75 |

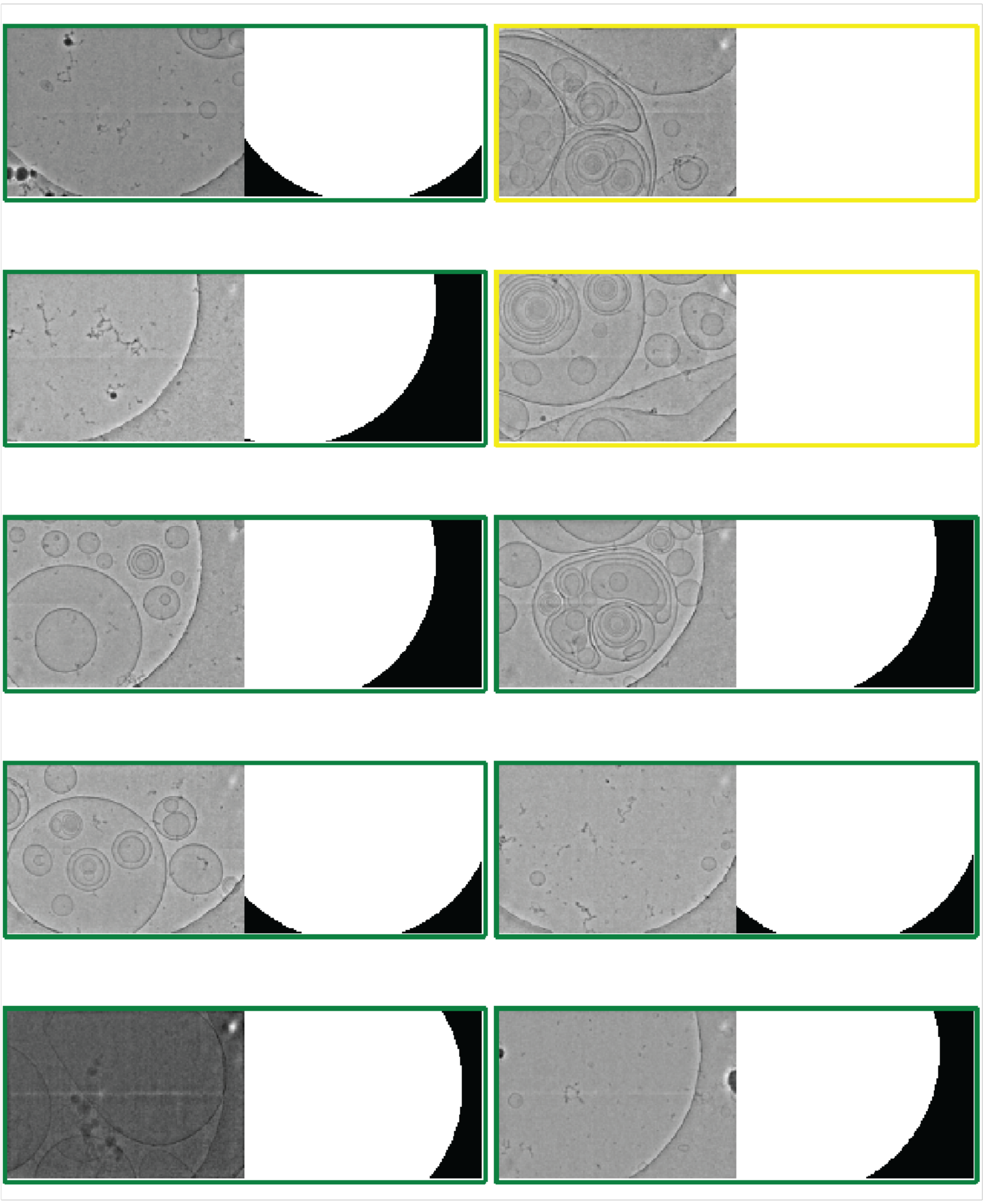

**Figure S1. CryoVIA detects foil holes and excludes the area for membrane segmentation, Related to Figure 2.**
Related to "Identification of membrane features". Example hole masking of 10 micrographs from the DOPC-DPPG data set using the edge detection GUI: micrograph (left) and identified binary mask (right). Green border: hole edge found, yellow border: no hole found.

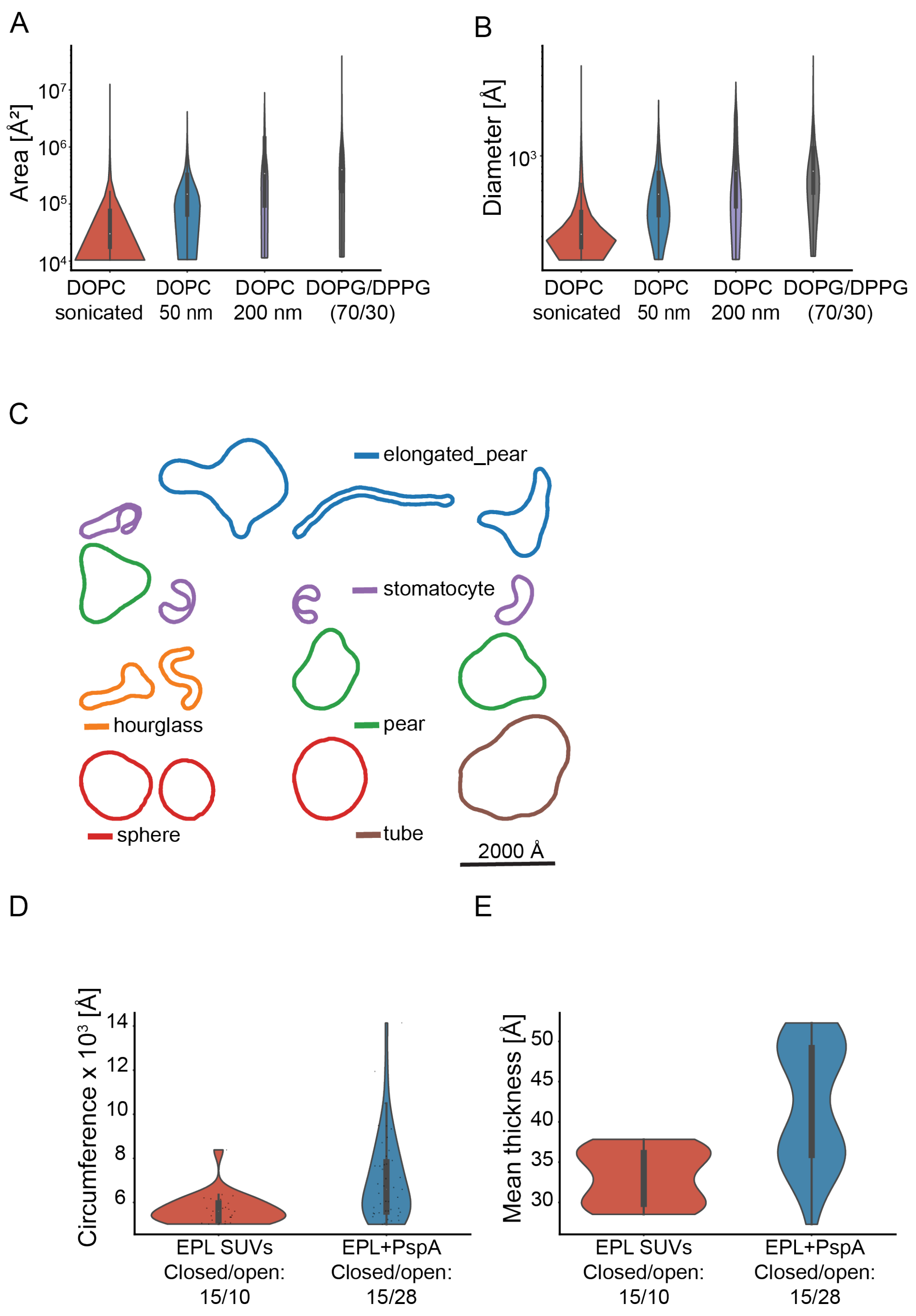

**Figure S2. CryoVIA evaluates structural features and analyzes shapes of membranes, Related to Figures 2, 5 and 6.**

(A, B) Violin plots of the evaluated vesicles from four data sets (DOPC sonicated (n = 38871), DOPC 50 nm size-filtered (n = 6061), DOPC 200 nm size-filtered (n = 1819), and DOPG/DPPG (70/30) (n = 2179)) with area (A) and diameter (B). (C) A total of 15 irregular shapes that have similarly high confidence score derived from the shape classifiers as in Figure 6D but were excluded due to irregular shapes that are not closely resembling any of the pretrained shapes. (D) Zoomed-in violin plot of the membrane length distribution of membranes with a length greater than 5000 Å of the EPL SUV and EPL + PspA data sets. (E) Zoomed-in violin plot of the mean thickness distribution of membranes with a length greater than 5000 Å of the EPL SUV and EPL + PspA datasets. (A, B, D, E) The white dot within each violin indicates the median, while the shaded area represents the kernel density estimate of the data distribution. The inner grey box represents the 25% to 75% interquartile range, while the vertical grey line includes values inside the interquartile range multiplied by 1.5. (D, E) The sample size is noted below the label.

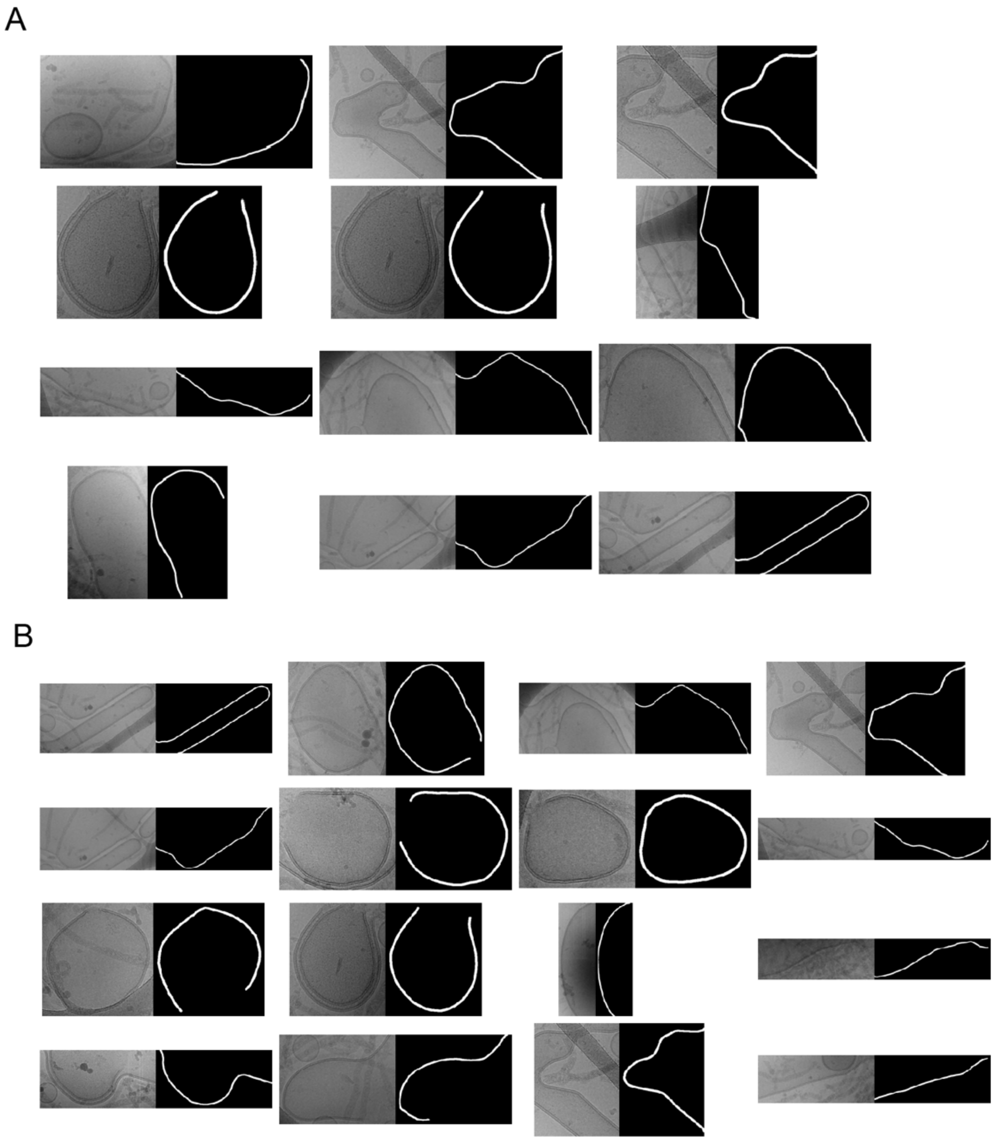

**Figure S3. CryoVIA's gallery of remodeled membranes and corresponding segmentation, Related to Figure 7.**
Gallery of EPL+PspA micrographs with membranes larger than the micrograph. (A) Some cropped non-closed membranes of the PspA data set with a length greater than 5000 Å. (B) Some cropped membranes of the PspA data set with a bilayer thickness greater than 45 Å.

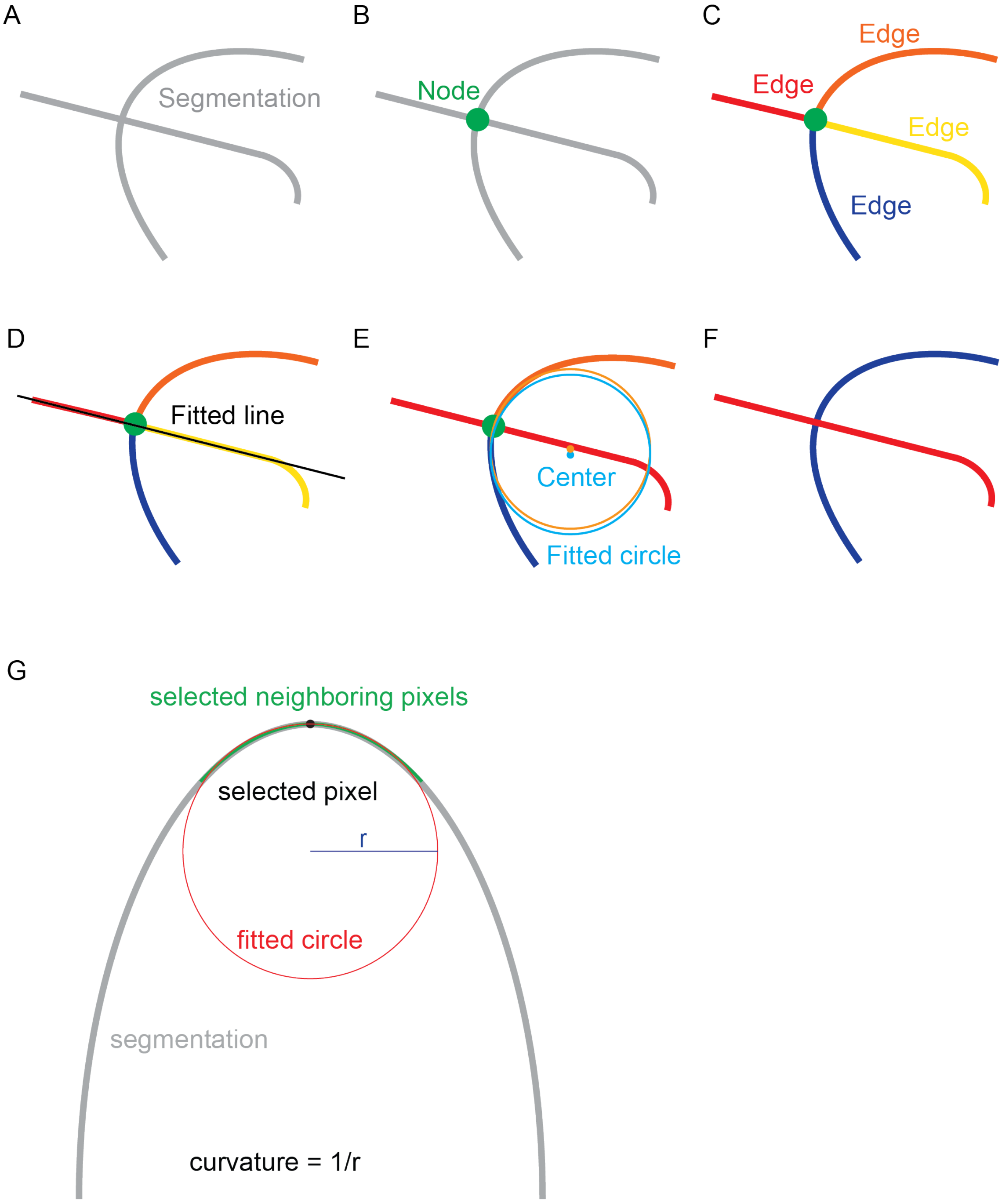

**Figure S4. CryoVIA's approach of resolving overlapping vesicles, Related to STAR Methods.**
(A) Example binary segmentation. (B) Identified node where vesicles overlap. (C) Four edges connected to the node. (D) Fitted a straight line through the red and yellow ends. (E) Combined red and yellow ends because the straight line fitted well close to the node resulting in two overlapping ends. Fitted circle for the orange and blue end close to the node and the center of the fitted circle. (F) Combined blue and orange edges to one membrane stretch as fitted circle centers were close to each other. (G) Illustration of the curvature estimation method. Grey: Segmentation. Black dot: Pixel location of curvature evaluation. Green: Neighboring pixel to fit a circle to. Red: Fitted circle. Blue: Radius of the fitted circle. The curvature is estimated as the reciprocal value of the radius of the best fitting circle.

A

B

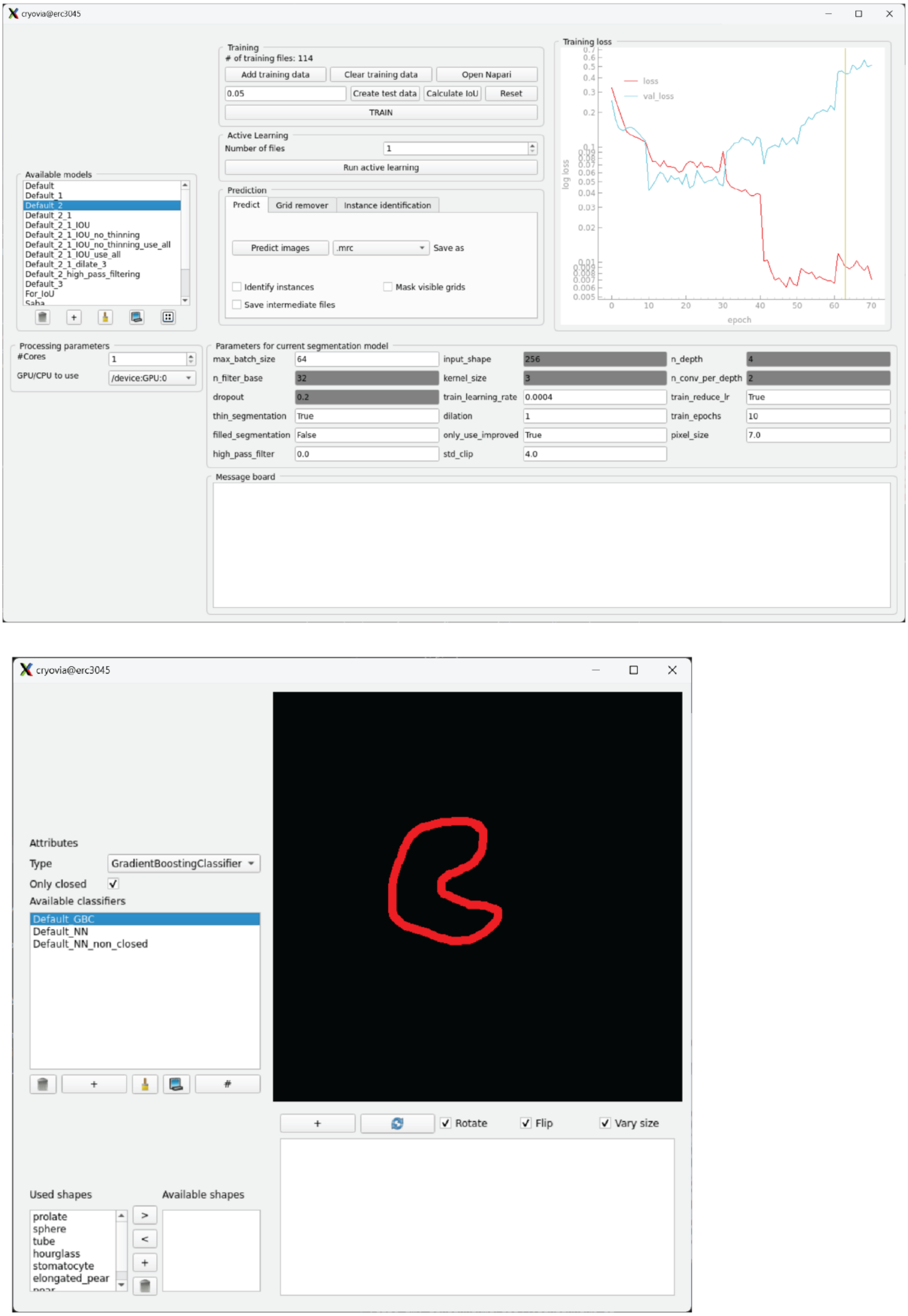

**Figure S5. Screenshots of CryoVIA GUI for segmentation, Related to STAR Methods.**
(A) Segmentation neural network training and modifier including parameters for segmentation model. (B) Display of results of shape classifier and modifier.

A

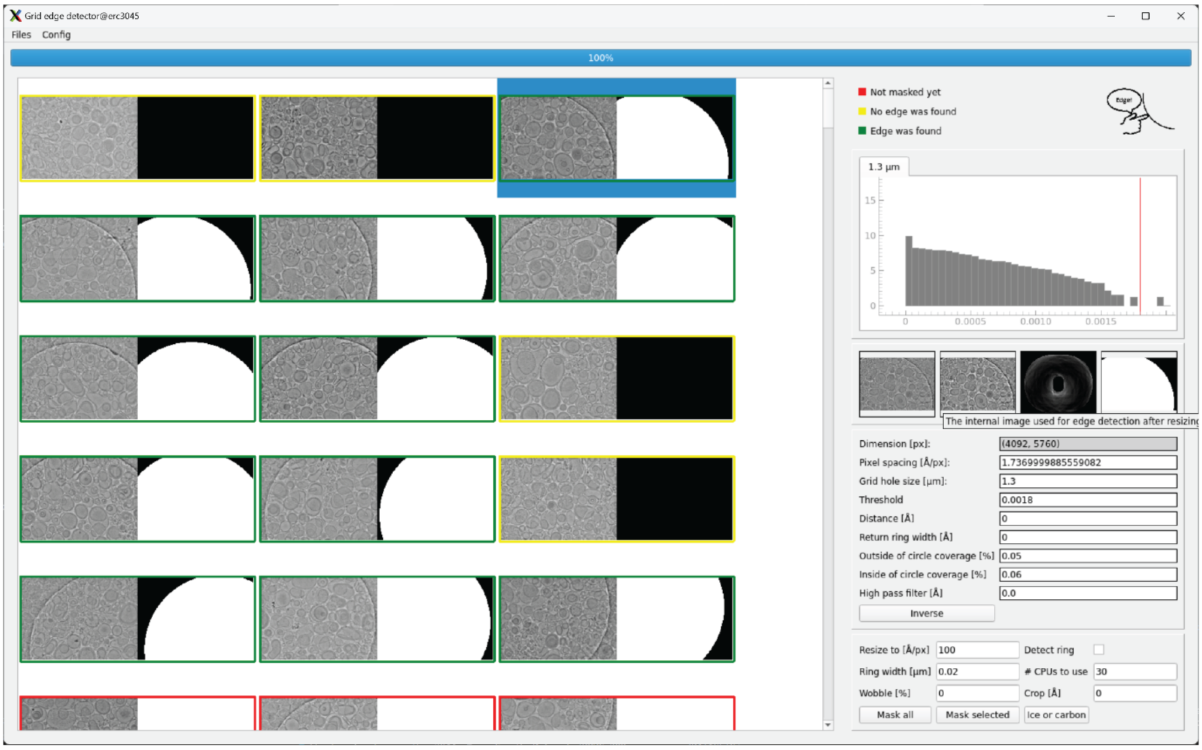

B

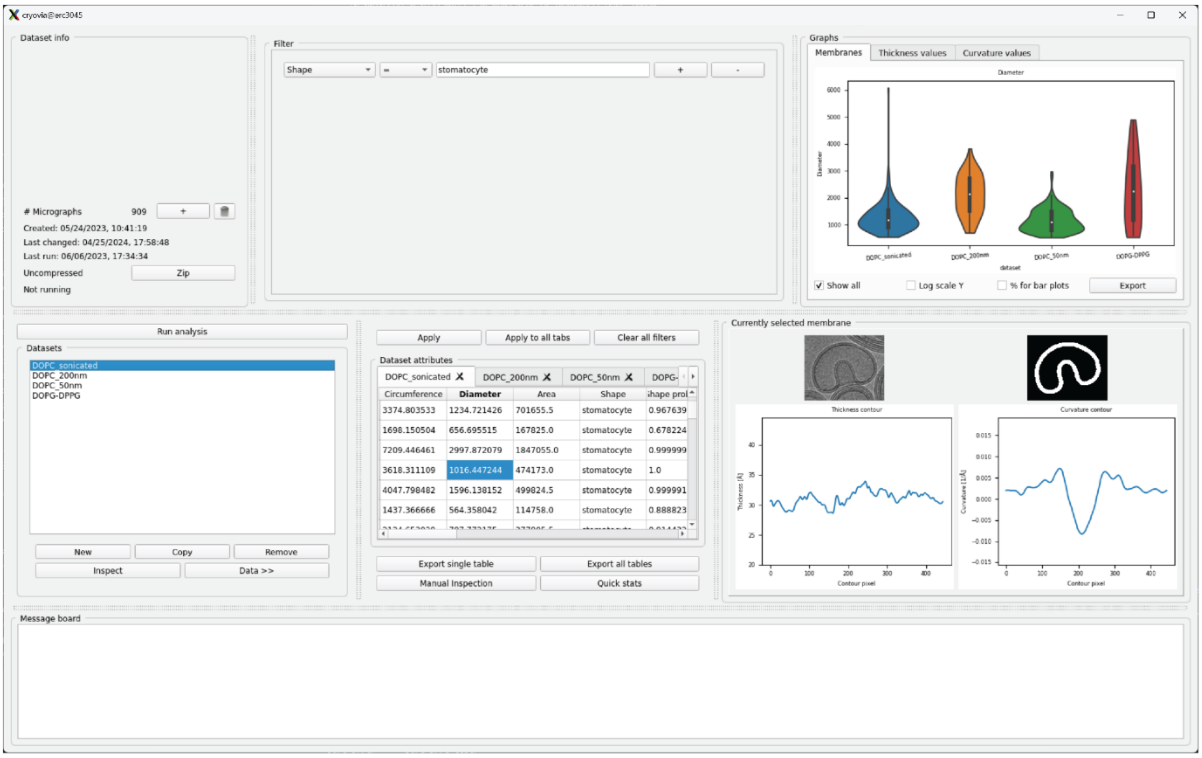

**Figure S6. Screenshots of CryoVIA GUI for analytical tools, Related to STAR Methods.**
(A) Foil hole edge detector. (B) Data analysis and data set creation/running.
